# Precise Transcript Reconstruction with End-Guided Assembly

**DOI:** 10.1101/2022.01.12.476004

**Authors:** Michael A. Schon, Stefan Lutzmayer, Falko Hofmann, Michael D. Nodine

## Abstract

Accurate annotation of transcript isoforms is crucial to understand gene functions, but automated methods for reconstructing full-length transcripts from RNA sequencing (RNA-seq) data remain imprecise. We developed Bookend, a software package for transcript assembly that incorporates data from different RNA-seq techniques, with a focus on identifying and utilizing RNA 5′ and 3′ ends. Through end-guided assembly with Bookend we demonstrate that correct modeling of transcript start and end sites is essential for precise transcript assembly. Furthermore, we discovered that utilization of end-labeled reads present in full-length single-cell RNA-seq (scRNA-seq) datasets dramatically improves the precision of transcript assembly in single cells. Finally, we show that hybrid assembly across short-read, long-read, and end-capture RNA-seq datasets from Arabidopsis, as well as meta-assembly of RNA-seq from single mouse embryonic stem cells (mESCs) can produce end-to-end transcript annotations of comparable quality to reference annotations in these model organisms.

**Summary statement:** Bookend is a generalized framework that utilizes RNA 5′ and 3′ end information hidden in RNA-seq datasets to accurately reconstruct transcriptomes including those from single cells.

## INTRODUCTION

The functions of genes depend on the amount and types of RNA molecules that they produce. Variation in transcript initiation, splicing and polyadenylation can generate an array of RNA isoforms, and cataloging how these RNA variants change across development and disease provides insights into corresponding gene functions^1–3^. Large-scale projects dedicated to the manual curation of gene annotations are extremely valuable, but are labor-intensive and thus limited in scope to the most well-studied organisms^4–7^. Moreover, multicellular organisms have difficult-to-access cell types that will inevitably be overlooked by even the most comprehensive annotation projects^8^. The completeness and accuracy of a reference annotation can considerably impact all downstream data analyses, from gene expression to predictions of gene function^9–11^. To understand how transcriptome architecture varies during development and in response to disease, it is therefore valuable to have an automated method that accurately identifies transcript isoforms. Accordingly, many computational tools have been developed for genome annotation including software that utilizes the massive and growing diversity of RNA sequencing (RNA-seq) technologies^12^.

A wide array of RNA-seq protocols have been developed to profile different aspects of the transcriptome, from strand-specific coverage of gene bodies^13^ to selective amplification of RNA 5′ ends^14–17^, 3′ ends^18,19^ or simultaneous capture of both ends^20,21^. Major recent advances have enabled the amplification of full-length transcripts from single cells^22,23^ or 3′ end capture from millions of cells^24–26^. In parallel, advances have been made for profiling RNA on “third-generation” long-read sequencing platforms such as PacBio and Oxford Nanopore single-molecule sequencers that can read a continuous DNA and/or RNA molecule many times the length of a typical transcript and yield end-to-end complete sequences of RNA molecules^27,28^.

Transcript assembly is the effort to distill information from RNA-seq experiments into a comprehensive annotation of the transcript isoforms present in the corresponding samples. Depending on the method, RNA-seq reads contain a broad spectrum of information content. At one extreme, single-end reads from a non-stranded RNA-seq protocol can be 50 nucleotides (nt) or shorter and sequenced from one end of a double-stranded cDNA fragment such that the resulting sequence is a random substring of an RNA molecule or its reverse complement. Paired-end reads contain two ends of a cDNA molecule and typically there is a gap of unknown length between the mate pairs. When aligned to a reference genome, paired reads may span more than one splice junction, indicating that these splicing events occurred in the same molecule. Some strand-specific RNA-seq protocols selectively sequence only first-strand or second-strand cDNA to preserve knowledge of the original mRNA molecule’s orientation^13^. Other protocols selectively capture and sequence a fragment immediately downstream of the RNA 5′ end or upstream of the 3′ end, demarcating precisely where that molecule begins or ends, respectively. Finally, the most information-rich reads come from long-read sequencing, in which the RNA or cDNA is read in its entirety without fragmentation. Long-read methods are a promising tool for transcript annotation, but current protocols are more error-prone per base sequenced, less sensitive, and more costly than comparable short-read experiments. Because the vast majority of existing RNA-seq data is in short-read format, nearly all assemblers have aimed to reconstruct transcripts from paired-end short reads. A long-recognized problem of assemblers is the inaccurate annotation of transcript start sites (TSS) and polyadenylation sites (PAS)^29,30^. Existing short-read assemblers infer TSSs and PASs at sharp changes in read coverage, but such changes can also be due to alignment errors, biased RNA fragmentation, sample degradation, or spurious intron retention. Long-read sequencing methods are designed to read RNA from TSS to PAS, but they remain susceptible to a variety of experimental artifacts^30^. The increasing adoption of long reads for transcript annotation has led to a separate suite of tools that summarize, collapse, or “polish” long reads to remove erroneous structures and present a set of representative isoforms from these reads^31,32^. For example, a recently developed transcript assembler reports the use of long reads in assembly by removing aligned segments with a high error rate and assembling the resulting gapped reads^33^. Transcript annotation would ideally integrate information from a variety of RNA-seq methods to determine the best evidence for transcript starts, ends and splicing patterns in a tissue-of-interest. However, current transcriptome assembly methods do not employ information about where RNAs begin and end. Here, we describe a method utilizing RNA 5′ and 3′ end information contained in RNA-seq datasets to accurately reconstruct transcriptomes including those from single cells.

## RESULTS

### A framework for end-guided transcript assembly

To determine whether RNA 5′ and 3′ end information can improve transcript assembly algorithms, we developed a generalized framework for identifying RNA ends in sequencing data and using this information to assemble transcript isoforms as paths through a network accounting for splice sites, transcription start sites (TSS) and polyadenylation sites (PAS). Because this software uses end information to guide transcript assembly, we named it Bookend. Importantly, Bookend takes RNA-seq reads from any method as input and after alignment to a reference genome, reads are stored in a lightweight end-labeled read (ELR) file format that records all RNA boundary features (5′ labels, splice donors, splice acceptors, gaps, 3′ labels), as well as the sample of origin for that read (see Supporting Notes). Assembly is then resolved at each locus with aligned reads through a four-step procedure (Fig1; see Methods and Supporting Notes). First, boundary labels from all aligned RNA-seq reads are clustered and filtered to demarcate a unique set of locus TSSs, PASs and splice junctions. Each locus is partitioned into a set of nonoverlapping “frags” defined as the spans between adjacent boundary labels. Four additional frags (S+, E+, S-, E-) denote the presence of a Start or End Tag on the forward or reverse strand. Second, a Membership Matrix is generated to redefine all aligned reads with respect to the locus frags. A read’s Membership includes each frag it overlaps and excludes each incompatible frag (e.g. a spanned intron, a region upstream of a TSS or downstream of a PAS). Reads with identical patterns of Membership are condensed to a single element (row) of the Membership Matrix, whose weight is the total coverage depth across the element by all reads of that pattern. Third, an Overlap Graph is constructed from the Membership Matrix elements and this directed graph is simplified by collapsing shorter elements into the elements that contain them. Finally, the Overlap Graph is iteratively traversed to resolve an optimal set of Greedy Paths from TSSs to PASs. These Paths describe a set of full-length transcript models best supported by the input reads. The Membership Matrix definition is flexible enough to utilize reads regardless of their length, alignment gaps, strand, or end information (FigS1B).

**Figure 1.**
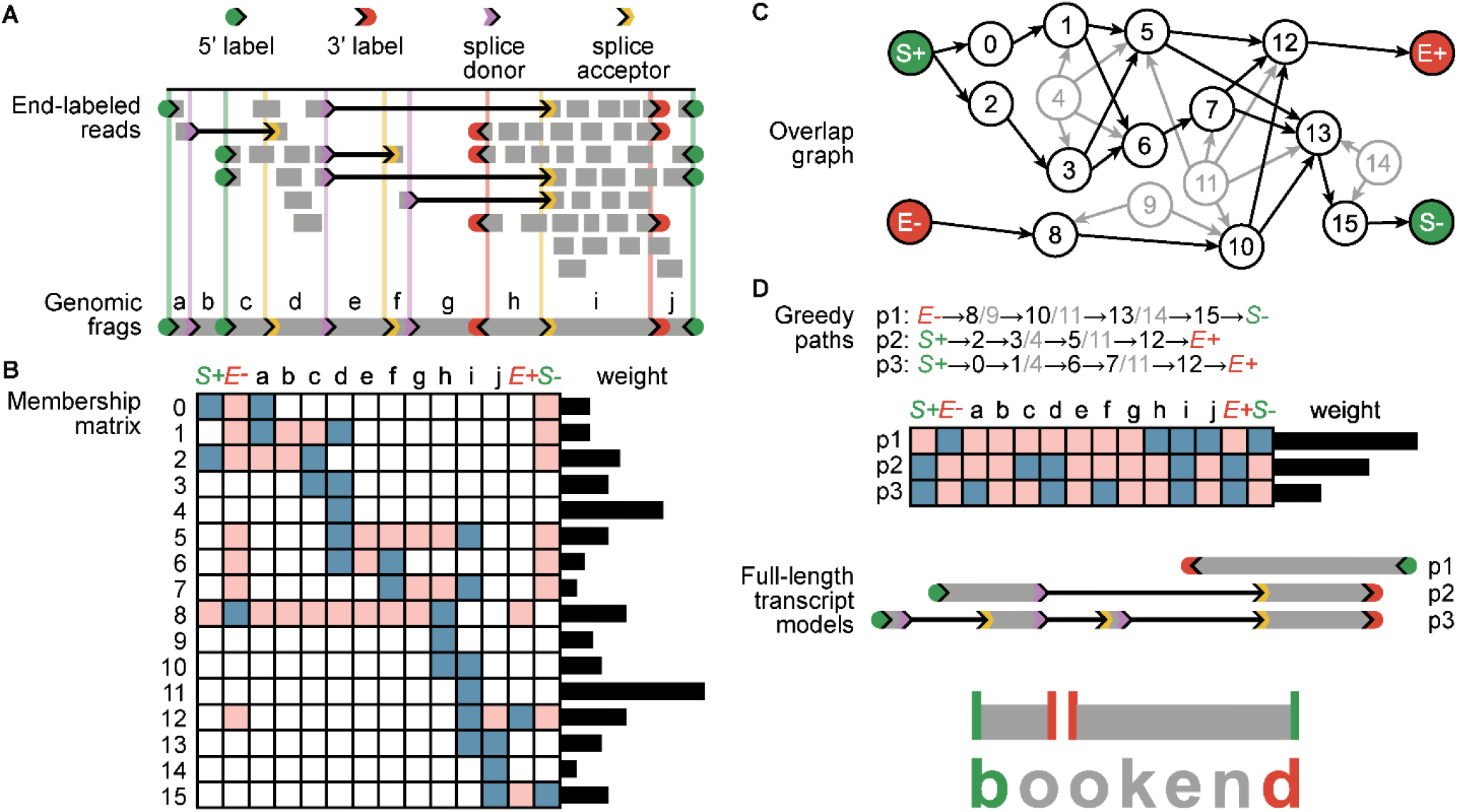
End-guided assembly with Bookend **(A)** Individual RNA-seq reads are mapped to a genome, recording which reads mark a transcript 5′ or 3′ end, and which reads span one or more splice junctions. Ranges between adjacent features are recorded as frags. **(B)** Each unique read structure is recorded in a condensed representation as one element in a Membership Matrix; blue-included, pink-excluded. The weight of each element is the coverage depth of matching reads (sequenced bases/length) across the element. **(C)** A directed graph is constructed between overlapping elements of the Membership matrix. Weights of contained elements (gray) are distributed proportionally to their containers. **(D)** A set of optimal paths through the graph is iteratively constructed from the heaviest unassigned elements. Complete Paths are output as full-length transcript annotations.

### End-labeled reads improve the quality of transcript assembly

Arabidopsis is an ideal model to benchmark transcript assembly in higher eukaryotes. The Arabidopsis genome is compact (∼119 megabases), contains few repetitive elements, and the TAIR10 reference annotation was extensively curated from expressed sequence tag (EST) data^7^. To determine whether assembly benefits from end-labeled reads, we examined libraries generated with the low-input sequencing method Smart-seq2 from Arabidopsis floral buds^16^. Two crucial steps in the Smart-seq2 protocol, template switching and preamplification, enrich for full-length cDNA with an oligo label at both the 5′ (template switching oligo, TSO) and 3′ (oligo-dT) end^22^. These oligos were trimmed from all reads and a record was kept of which end label was found (5′, 3′, or no label) before mapping to the genome. As anticipated, a small percentage of reads were found with either label (Fig 2A; Supplemental Table 1). All reads were aligned to the Arabidopsis genome, and the terminal positions of 5′- and 3′-labeled reads were retained as “Start Tags” and “End Tags”, respectively. Of End Tags mapping to annotated genes, 88% mapped near PASs, defined as the last decile of the gene or up to 100nt downstream (Fig2B). Start Tags had lower specificity for TSSs, with only 48% of Start Tags in the first decile of genes or up to 100nt upstream. Template switching is known to readily occur at RNA 5′ ends derived from in vivo or in vitro RNA decay. However, a subset of reads contain an intervening G between the TSO and the genome-aligned sequence, indicating a 7-methylguanosine cap on the template RNA^16,34,35^. The upstream untemplated G (uuG)-containing Start Tags were classified as Cap Tags. Cap Tags were rare relative to all Start Tags (9%), but were much more specific to TSSs with an average of 88% of Cap Tags within each gene mapping near the 5’ end (Fig2B). To optimize detection of true transcript 5’ and 3’ ends, the Tag Clustering algorithm designed for Bookend defines Tag weight as a function of total read depth and applies a bonus to Cap Tags over non-uuG Start Tags (See Supplemental Note: “Tag Clustering”).

**Figure 2.**
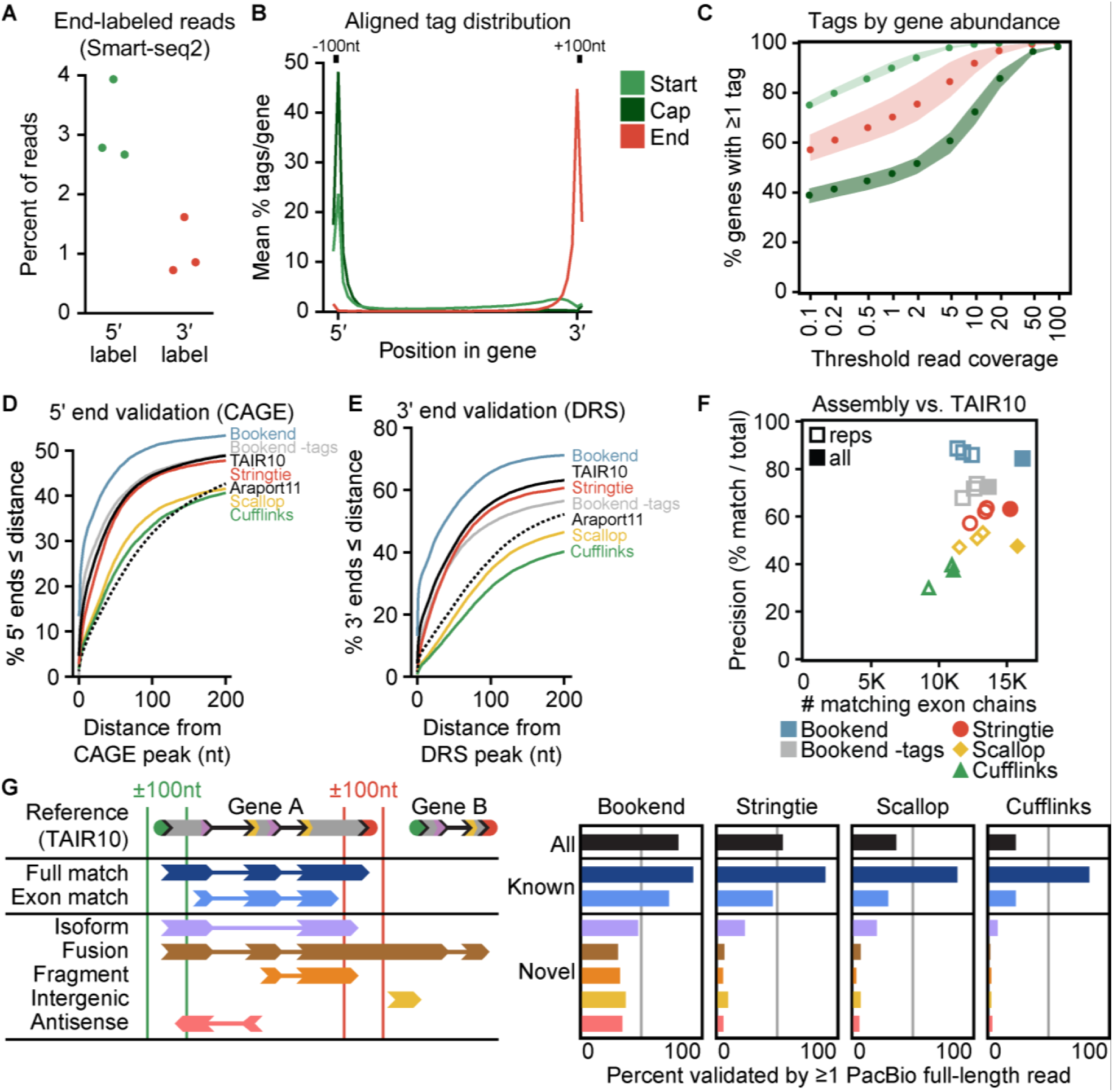
End-labeled Smart-seq2 reads accurately detect transcript 5′ and 3′ ends. **(A)** Percent of reads in three Smart-seq2 libraries that contained a 5′-labeled or 3′-labeled junction, respectively. **(B)** Average signal strength per gene of Start, End, and Cap Tags along gene bodies in 50 bins with an additional 100nt flanking each gene boundary. Start Tag, any 5′ label; Cap Tag, 5′ label with upstream untemplated G (uuG); End Tag, 3′ label. **(C)** Likelihood of a gene to possess ≥1 Start, Cap, or End Tag as a function of aligned read coverage (average read depth/base). **(D)** Cumulative frequency of annotated 5′ ends as a function of distance from the closest CAGE peak36. **(E)** Distance of 3′ ends from the nearest DRS peak37 as in (D). **(F)** Performance of three transcript assemblers, measured by total number of reference-matching exon chains (x-axis) vs. percent of assembled transcripts that match the reference (y-axis). **(G)** (Left) Schematic depicting classifications of assembled transcripts against the closest TAIR10 reference isoform. (Right) Rate of validation by PacBio full-length non-chimeric (FLNC) reads for different assemblies, grouped by classification.

Despite end-labeled reads being relatively rare, the preamplification process should ensure that a TSO or oligo-dT sequence is at each end of every cDNA molecule prior to tagmentation. Therefore, we expected end-labeled reads to be distributed widely across the genome wherever reads exist. As predicted, the majority of genes with >0 read coverage contained ≥1 Start Tag and ≥1 End Tag, and the likelihood of finding a Start or End Tag increased as a function of total read coverage (Fig2C). Of all genes with at least 1x, 10x and 100x read coverage, 73.3%, 94.4% and 99.2% possessed both a Start and End Tag, respectively.

To assess whether end-labeled reads mark real TSSs and PASs at nucleotide precision, Bookend was used to assemble all floral bud Smart-seq2 reads either with or without utilizing Start and End Tags. Additionally, three leading short-read transcript assemblers were used with comparable settings (see Methods): StringTie2^33,38^, Scallop^39^, and Cufflinks^40^. Publicly available Arabidopsis CAGE^36^ and Direct RNA-seq (DRS^37^) datasets were used to validate 5′ and 3′ ends, respectively. All three of these widely-used assemblers output thousands of single-exon unstranded fragments, which were ambiguous with regard to which end is 5’ or 3’ and thus were discarded from further analyses (Supplemental Table 2). Bookend-defined TSSs based on Start/Cap Tags were more likely to have a CAGE peak within 200nt than 5′ ends reported either by Bookend without the use of Start Tags, the three leading assemblers, or even the current Arabidopsis reference annotations (Fig2D). Likewise, a higher proportion of Bookend-identified PASs were supported by DRS reads than PASs reported by the other transcript assemblers and Arabidopsis reference annotations (Fig2E). At the nucleotide level, Bookend-defined transcript boundaries were more than twice as likely to agree with the exact experimentally-determined TSS and PAS peak positions than the most accurate reference annotation (TAIR10), while the other three assemblers reported transcript boundaries less accurate than TAIR10 (FigS2A-B). Strikingly, even the Bookend 5′ and 3′ ends >100nt from any reference still possessed known sequence motifs associated with TSS and PAS, respectively, whereas sequence content around novel ends from Cufflinks, Scallop, and StringTie2 is largely incoherent (FigS2C-D). In addition to a dramatic increase in transcript boundary accuracy, 16,158 exon chains predicted by Bookend fully matched a TAIR10 reference transcript, which was higher than when end-labeled reads were ignored (13,660) and exceeded the totals from Scallop (15,785), StringTie2 (15,253) or Cufflinks (11,051) (Fig2F). Therefore, Bookend correctly builds more known transcripts than other assemblers and Bookend-annotated 5′ and 3′ ends were more precise than even the most accurate Arabidopsis reference annotation.

In addition to known transcripts, Bookend constructed 2,979 isoforms not present in TAIR10, which was 66% fewer than StringTie2 (8,886), 83% fewer than Scallop (17,400), and 84% fewer than Cufflinks (18,934). An assembled transcript may fail to match TAIR10 either because the assembly is incorrect or because the reference is incomplete. To distinguish between these possibilities, two long-read SMRT cells of floral bud RNA were sequenced with the PacBio platform to yield 547,910 full-length non-chimeric (FLNC) reads. All short-read assemblies were partitioned into 7 different classifications based on their relationship to the most similar TAIR10 model (Fig2G). A transcript model was considered experimentally validated if at least one aligned PacBio read fully matched the model (entire exon chain, ±100nt ends). Of all Bookend transcripts, 81.2% were supported by PacBio data, which surpassed the validation of transcripts predicted by StringTie2 (54.7%), Scallop (35.9%) or Cufflinks (22.3%) (Fig2G; Supplemental Table 2). Reference-matching transcripts have a higher average estimated abundance than non-reference transcripts, making the latter more difficult to validate with the limited throughput of long-read sequencing (FigS2E). Despite this limitation, 42.3% of non-reference Bookend assemblies were fully supported by at least one PacBio read, which was substantially higher than the validation rate of non-reference transcript assemblies generated by StringTie2 (15.9%), Scallop (11.6%), and Cufflinks (4.3%) (Fig. 2G). Taken together, these results demonstrate that end-guided assembly using latent RNA end information enables precise transcript reconstruction from short-read datasets.

### Hybrid assembly refines and complements long-read RNA-seq

Long-read sequencing technologies do not obviate the need for transcript reconstruction. Various sources of technical and biological noise result in fragmented or improperly spliced long reads^30,41^. Long-read approaches also suffer from a higher base-level error rate compared to short-read platforms^42^. Error correcting methods such as Circular Consensus Sequencing (CCS) require reverse transcription and cDNA amplification, which are susceptible to mispriming and template-switching artifacts^43,44^. This has driven the ongoing development of tools to refine transcript models derived from long reads^31,32^. Additionally, StringTie2 was recently repurposed to assemble long reads^33^.

To quantify potential sources of error, PacBio FLNC reads were aligned to the genome and processed by the Bookend pipeline to identify and remove template-switching artifacts, oligo-d(T) mispriming events at A-rich regions, and exons with a high alignment error (Fig3A). Across both SMRT cells, 95.4% of reads aligned successfully, and 97.0% of alignments did not contain any high-error exons, defined as the total length of mismatches, inserts, and deletions exceeding 10% of the exon length. However, 14.1% of all FLNC 3′ end labels were removed due to alignment failure or the presence of an A-rich region immediately downstream of the oligo-d(T) junction. If treated as genuine 3′ ends, these reads can cause false annotation of 3′-UTRs or putative transcripts antisense or intergenic to known genes^43^ (FigS3A). Direct RNA sequencing bypasses oligo-d(T) priming and was used to produce a map of genuine Arabidopsis PAS^37^. These sites show a distinct pattern of nucleotide enrichment, including a C/A dinucleotide motif at the cleavage and polyadenylation site itself, and a U-rich upstream element (USE) and downstream element (DSE) (Fig3B). Three tools were used to reduce the PacBio FLNC data into a unique set of transcripts: the Iso-seq3 clustering algorithm from PacBio, assembly by StringTie2, and end-guided assembly by Bookend. All 3 methods could recapitulate known PAS motifs at the set of 3′ ends within 100nt of a TAIR10-annotated PAS. In contrast to Bookend, StringTie2-annotated 3′ ends showed a slight A-richness at novel 3′ ends, and both Iso-seq3 and StringTie2 annotations contain thousands of putative novel antisense or intergenic RNAs whose 3′ ends are extremely A-rich (Fig3C). Therefore, Bookend retains genuine novel PAS by filtering against known 3′ artifacts.

**Figure 3.**
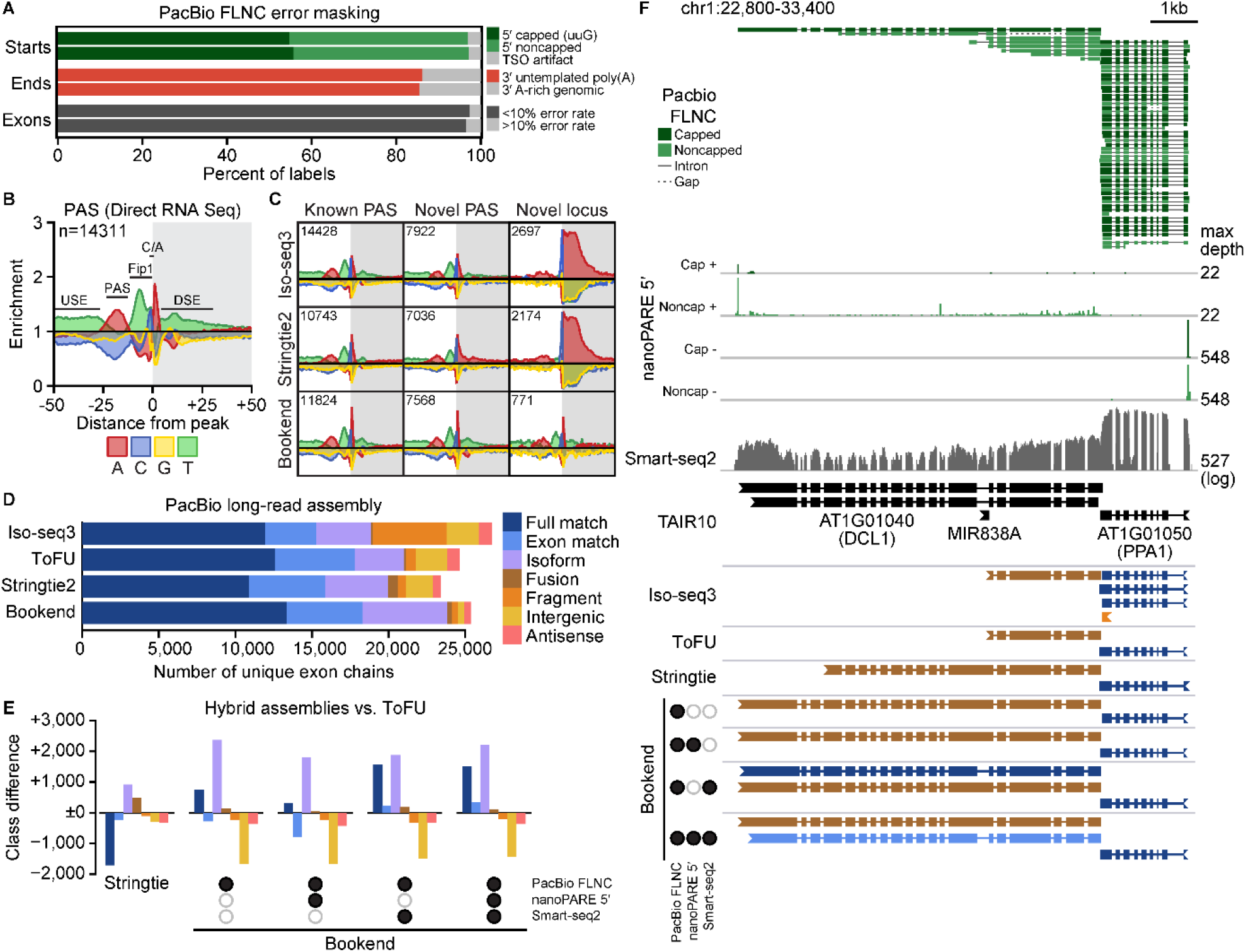
Long-read sequencing is augmented by hybrid assembly **(A)** Artifacts identified in PacBio FLNC reads from two SMRT cells by alignment to the Arabidopsis reference genome. **(B)** Nucleotide frequency enrichment in a ±50nt window around poly(A) sites (PAS) identified by Direct RNA Seq37. **(C)** Nucleotide enrichment around 3′ ends of transcripts constructed from PacBio reads by Iso-seq3 (top), StringTie2 (middle), and Bookend (bottom) at sites overlapping a TAIR10 PAS (left), novel PAS at a known gene (middle), and novel antisense or intergenic loci (right); colors and scales as in **B. (D)** Classification against TAIR10 of transcripts constructed by four long-read strategies: Iso-seq3 clustering, cluster collapse by ToFU, and FLNC assembly by StringTie2 and Bookend. **(E)** Effect of long-read assembly on the number of transcripts by class (colored as in **D**) by StringTie2 (left) or hybrid assembly with one or more tissue-matched sequencing libraries by Bookend (right). Bars show difference vs. ToFU-collapsed Iso-seq3 clusters. **(F)** Integrative Genomics Viewer (IGV) image of the Arabidopsis *DICER-LIKE1* (*DCL1*) locus. From top to bottom: PacBio reads colored by 5′ end label, nanoPARE capped and noncapped read 5′ end frequency, Smart-seq2 read coverage, TAIR10 reference models, and long-read assemblies colored by classification vs. TAIR10 as in **D**.

Another major source of transcript assembly error is truncated 5′ ends due to premature template switching during reverse transcription or amplification of degraded RNA. Although 50% of FLNC alignments matched a full-length TAIR10 transcript, most were copies of a few highly-expressed genes. After collapsing alignments into sets of unique exon chains, full-length reference transcripts accounted for only 18% of all unique chains, and 25% of unique chains were fragments of known TAIR10 transcripts missing one or more exons (Supplemental Table 3). Clustering by Iso-seq3 removes some fragments, and they can be further reduced after alignment by collapsing 5′ truncations with Transcript isOforms: Full-length and Unassembled (ToFU) ^45^ (Fig3D). However, it was unknown whether an assembly algorithm would further improve the quality of long-read annotations. Surprisingly, passing the FLNC data through StringTie2 yielded 1,704 fewer full-length reference matches compared to ToFU, and the number of transcripts classified as fusions of two different genes increased nearly four-fold (Fig3D-E). Because the Arabidopsis genome is compact with an average of only 1.5 kilobases (kb) between adjacent genes, assembly algorithms agnostic to 5′ and 3′ end information risk mis-annotating fused genes due to spurious read-through transcripts (FigS3A). By contrast, end-guided assembly of PacBio FLNCs with Bookend yielded 761 more full-length reference matches than ToFU, fewer than half as many fusions as StringTie2, and over a thousand more putative novel isoforms than both (Fig3D, Table S3).

Bookend’s assembly model is general enough to mix reads from different sequencing strategies. Therefore, we generated “hybrid assemblies” from combinations of PacBio FLNCs with Smart-seq2 and/or nanoPARE (a 5’ end sequencing strategy) from floral bud RNA^16^. All hybrid assemblies had higher precision than assembling long reads alone, and up to 809 more full-length matches could be identified (Fig3E, FigS3C, Supplemental Table 3). For example, *DICER-LIKE1* (*DCL1*) encodes the Arabidopsis *Dicer* homolog required for microRNA biogenesis, and its mRNA is maintained at low cellular abundance through an autoregulatory negative feedback loop involving two microRNAs, miR162 and miR838, the latter of which is encoded in intron 14 of its own gene^46,47^. Long reads alone were not sufficient to define the canonical 6.2 kilobase *DCL1* transcript because 7 of 8 PacBio reads mapping to *DCL1* were non-capped truncations, and intron 14 was retained in the only full-length read (Fig3F). By synthesizing information from multiple modes of sequencing, hybrid assembly with Bookend built a more complete transcript catalog that includes both the fully-spliced isoform and the isoform that retains MIR838. As a final refinement, a hybrid assembly that requires the presence of Cap Tags at transcript 5′ ends yielded a transcriptome with a 74.6% global concordance with the TAIR10 annotation. We report this hybrid assembly of long, short and 5′ end reads as the Bookend Floral Bud annotation (Supplemental Dataset 1-2).

### Transcript discovery from single-cell sequencing

Bookend achieved comparable precision assembling Arabidopsis transcriptomes from either long reads or short reads generated by Smart-seq2, which is a protocol routinely used for single-cell RNA sequencing (scRNA-seq) (FigS3C). However, scRNA-seq poses multiple hurdles to accurate assembly. Amplifying the few picograms of RNA in a single cell exacerbates biases and artifacts during reverse transcription^22^, and dropouts from inefficient RNA capture place limits on accurate isoform quantification from scRNA-seq^48^. Additionally, scRNA-seq has been most widely adopted in the study of mammalian systems. The mouse genome (and likewise the human genome) is roughly 30 times larger than the Arabidopsis genome with an average of twice as many introns per gene and nearly three times the number of annotated isoforms. Additionally, mouse introns can exceed 100kb and are on average 36 times longer than in Arabidopsis. Many isoforms per gene and large spans of non-genic sequence make it considerably more challenging both to assemble transcripts and to validate which assemblies are correct. To evaluate Bookend’s utility on mammalian scRNA-seq data, we tested it on a dataset designed for single-cell benchmarking^49^ which contains a set of synthetic Spike-In RNA Variants (SIRVs) added prior to cell lysis. SIRVs were designed to present a challenge to isoform quantification tools by mimicking complex mammalian genes^50^. The 69 synthetic transcripts map to 7 regions on a hypothetical genome in a way that recapitulates canonical and non-canonical splicing variation, antisense transcription and alternative 5′ and 3′ ends with up to 18 isoforms per gene (Fig S4A). SIRV Mix E2 contains molecules in four discrete concentrations so that each locus has major and minor isoforms that vary in relative abundance by up to 128-fold. SMARTer library preparations from 96 single mouse embryonic stem cells (mESCs) were deeply sequenced, with an average of 7 million aligned paired-end 100bp reads per cell (Supplemental Table 4) including an average of just over 500,000 SIRV-mapping reads per cell. Bookend correctly reconstructed (full splice match and ≤100nt error on both ends) an average of 22.6 transcripts per cell, which was higher than either Scallop (16.3) or StringTie2 (13) (Fig5AB). Moreover, Bookend assembled fewer false SIRVs than StringTie2 and especially Scallop (Fig5B). To test a relationship between performance and sequencing depth, cells were progressively combined into pairs, then sets of 4, 16, 32, and a full merge of reads from all 96 cells. The relative performance of the three assemblers was stable over two orders of magnitude of input with the F-measure (harmonic mean of precision and recall) slightly rising for Bookend as the sequencing depth increased and slightly decreasing for the others (Fig5B). Importantly, Bookend consistently assigned a higher estimated abundance to true transcripts, and false assemblies were more concentrated in the low abundance regime than for other assemblers (Fig5A). Overall precision on SIRVs averaged 55.9% for Bookend (vs. 39.6% StringTie2, 22.5% Scallop), and precision on the most abundant half of assemblies was 74.2% (vs. 48.2% StringTie2, 28.4% Scallop).

End-labeled reads mapping to the mouse genome were also assembled for each cell, and transcript models were compared to RefSeq mm39. All matching exon chains were considered matches, and precision was measured as the percent of all assemblies that match RefSeq. Recall was defined by tallying all transcripts correctly assembled at least once and counting the proportion of this transcript set found per cell. Although recall was considerably worse for Bookend (average 7.9%) than other methods (StringTie2 16.6%, Scallop 16.5%), precision was multiple times higher (76.3% Bookend, 29.0% StringTie2, 26.5% Scallop; FigS4B). Assemblies were repeated for two replicates of Smart-seq2 data from the same experiment with comparable results demonstrating that end-guided assembly is consistent across full-length sequencing protocols (FigS4B).

As with TAIR10, RefSeq is almost certainly incomplete, and non-reference-matching assemblies could still be valid. To experimentally validate non-RefSeq mESC assemblies, three validation datasets were used: uuG-containing SLIC-CAGE^17^ reads from mESCs for 5’ end validation, mESC 3P-Seq^51^ reads for 3’ end validation, and a database of long noncoding RNAs identified by intergenic Capture Long-read Sequencing (CLS^52^) for full-length validation of novel intergenic loci. An assembly was considered validated by a method if at least one read directly supported an assembled transcript’s respective structure(s). Assemblies with 5’ ends ≤100nt away from a RefSeq TSS contained “known” TSSs, and all others possessed “novel” TSSs. Likewise, assemblies with 3’ ends ≤100nt from their matching reference polyadenylation sites were considered “known” PASs and all others were “novel”. An average of 99.7% of Bookend, 83.9% of Scallop and 79.0% of Stringtie2 single-cell assemblies with a known TSS had at least one SLIC-CAGE read within 100nt (Fig4C). Moreover, the majority of novel, antisense and intergenic TSSs from Bookend transcripts were supported by at least 1 capped SLIC-CAGE read, whereas no novel group from StringTie2 or Scallop surpassed a 25% validation rate. The 3P-Seq dataset had fewer total reads and was less sensitive overall, but it still supported 19.9% of intergenic Bookend assembly 3’ ends, compared to 1.4% for Scallop and 0.8% for StringTie2 (Fig4D). By comparing against the CLS atlas we could validate the full structure of intergenic mESC assemblies. Bookend assembled a very small number of novel intergenic transcripts per cell (average 33 vs. 1209 by StringTie2 and 1073 by Scallop), but 49% of these were supported by one or more reads from the CLS atlas, compared to just 3% for Scallop intergenic assemblies and 0.3% for StringTie2 (Fig4E). Finally, because Cap and End Tags were extremely sparse in each cell (Supplemental Table 4), we hypothesized that the lower sensitivity could be explained by dropout of end labels. Supplying the mESC SLIC-CAGE (5’) and 3P-seq (3’) datasets to a Bookend hybrid assembly raised recall from 7.9% to 18.2% and retained a precision of 67.2% (FigS4B). Therefore, end-guided assembly of single-cell RNA-seq data can be used to identify genuine transcriptional novelty that is otherwise masked by noise.

**Figure 4.**
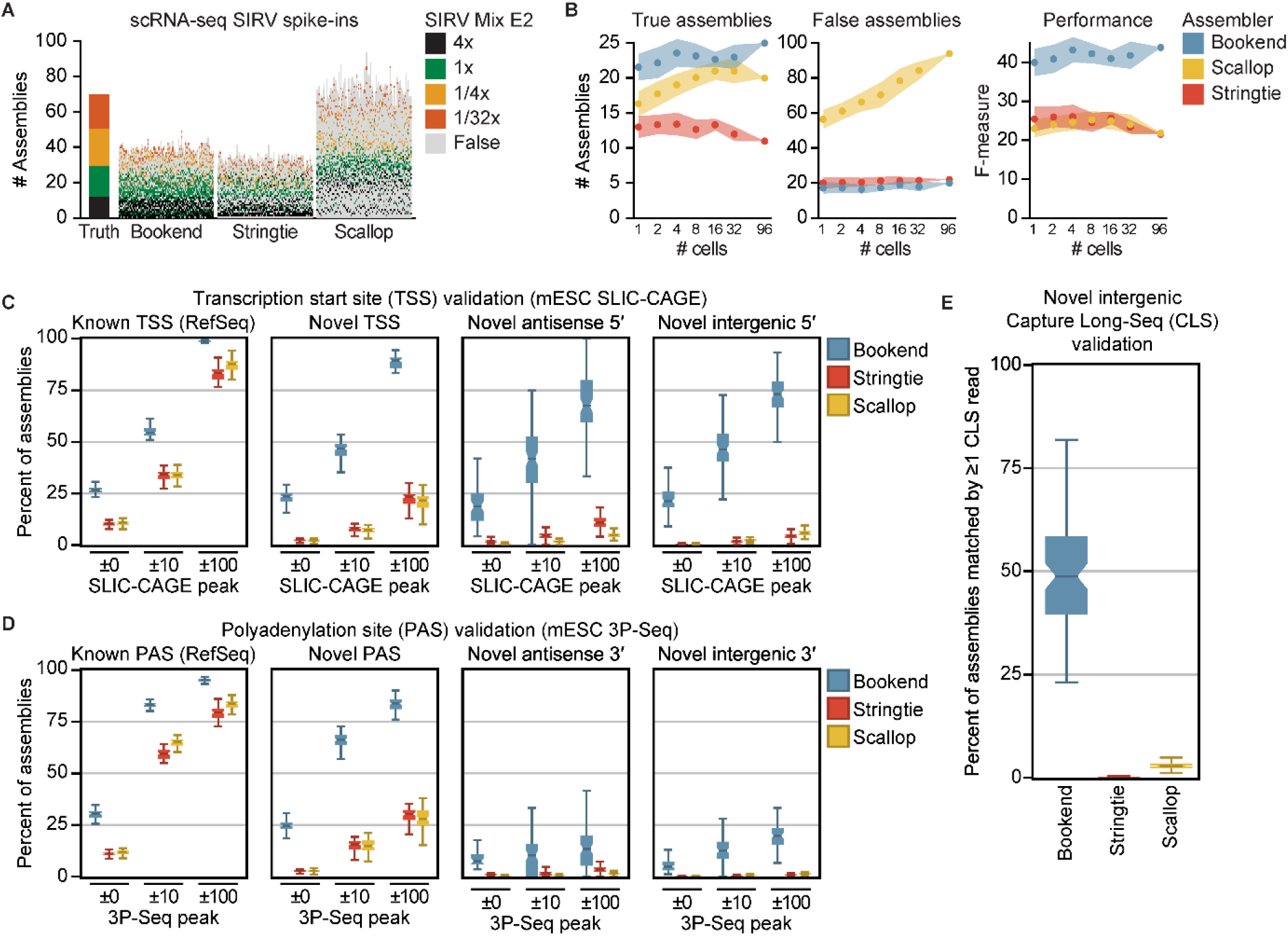
Bookend performance on single mouse cells **(A)** Reconstruction of Spike-In RNA Variants (SIRVs) from 96 paired-end 100bp SMARTer libraries of single mESCs. Each vertical bar depicts the assemblies from one cell, ordered from highest (bottom) to lowest (top) estimated abundance. Colored boxes match a true isoform of the given input concentration; gray boxes are false assemblies. **(B)** SIRV assembly performance as a function of increasing sequencing depth. F1 score (right) is the harmonic mean of sensitivity and precision. **(C)** Boxplots showing percent validation of 5′ ends with SLIC-CAGE support within the given windows for 96 single mESC assemblies. **(D)** Boxplots as in **C** showing 3′ end validation by 3P-Seq peaks. **(E)** Percent of intergenic assemblies (no overlap with RefSeq) in single cells which have ≥1 matching Capture Long-Seq read from the mouse CLS atlas.

### Condensed assembly and meta-assembly

A defining feature of single-cell experiments is that many individual cells are profiled in parallel. While sensitivity in an individual cell is low, information across multiple cells can be combined to achieve a more complete view of the experiment. Tools have been developed for transcript “meta-assembly” of reads from multiple sources. By modeling for variation across samples, meta-assemblers achieve higher precision than standard assembly on the same set of reads^53,54^. To measure the impact of meta-assembly, a series of assemblies on subsamples of all 706 million aligned single-cell mESC reads was first performed with StringTie2 and Scallop, as well as Bookend with and without the addition of mESC SLIC-CAGE and 3P-seq libraries (Fig5A). The mean number of reference-matching transcripts varied greatly across assemblers on single cells (1,656 Bookend, 3,711 Bookend hybrid, 2,904 StringTie2, 2,831 Scallop), but the magnitude of difference decreased with progressive doublings, up to the full set of 96 cells (12,794 Bookend, 13,762 Bookend hybrid, 13,524 StringTie2, 15,611 Scallop). By contrast, non-matches grew linearly with input. Bookend consistently assembled roughly an order of magnitude fewer non-matching transcripts than other assemblers across all input levels. From the full 96-cell dataset Scallop identified the most matches, but this was dwarfed by nearly 13 times the number of assemblies that failed to match RefSeq (201,631 Scallop, 100,646 StringTie2, 14,301 Bookend, 15,711 Bookend hybrid). By assuming non-matches to be mostly false, we calculated recall and precision as before and combined them to track the relationship between overall performance (F-measure) and input. F-measure of Bookend and Bookend hybrid assembly continued to improve with increasing input, but Scallop and StringTie2 began to decline above 4 and 16 cells, respectively, due to the growth of non-matches outpacing matches (Fig 5B). Consistent with previous reports, we see that standard assemblers suffer from an input-dependent decay in precision^53,54^.

**Figure 5.**
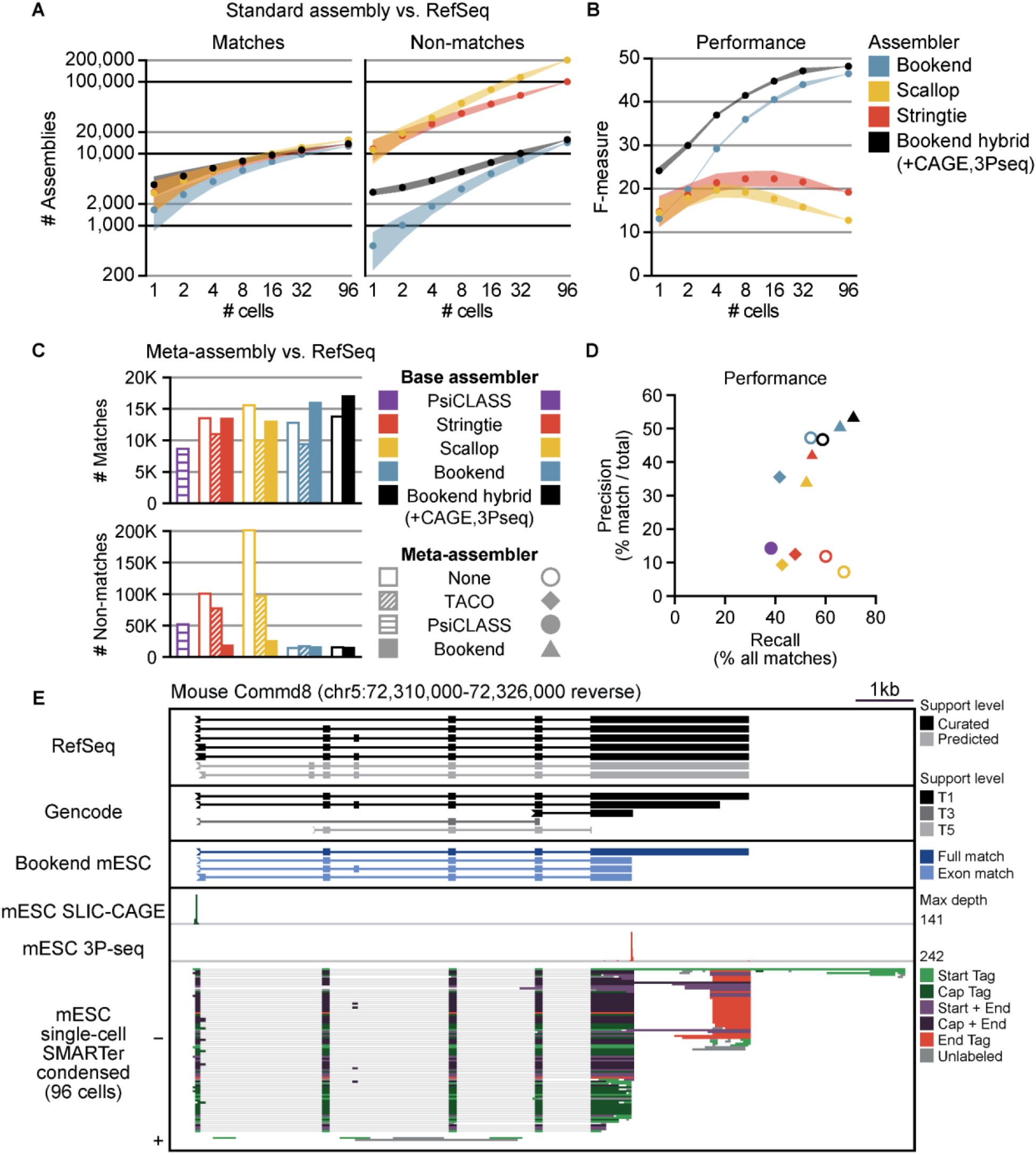
End-guided meta-assembly accurately integrates single cell data **(A)** Performance of assemblers with input from increasing numbers of single mESC cells. Assemblies with a matching exon chain to a RefSeq transcript (left) or no match to a RefSeq transcript (right). **(B)** F-measure of assemblies, where recall is the proportion of all transcripts assembled by ≥1 strategy and precision is matches/total assemblies. **(C)** Comparison of Bookend meta-assembly to standard assembly and other meta-assemblers. Number of RefSeq-matching transcripts assembled (top) or the number of non-matches (bottom). **(D)** Precision/recall plot of the 12 assemblies from **C**; recall and precision calculated as in **B. (E)** IGV browser image of the Commd8 gene. From top to bottom: RefSeq, Gencode, and Bookend mESC annotations, 5′ ends from mESC SLIC-CAGE, 3′ ends from mESC 3P-seq, Bookend-condensed partial assemblies from 96 single mESCs.

As an alternative approach, two published meta-assemblers were used to process the 96-cell dataset. TACO builds a consensus annotation by re-defining transcript boundaries through “change-point detection” on a set of files from any standard assembler^53^, whereas PsiCLASS generates the individual assemblies and performs meta-assembly through a consensus voting system^54^. The flexibility of Bookend’s framework allows its assembly algorithm to be run on assemblies, including its own output. To test the efficacy of meta-assembly with Bookend, each of the 96 mESC cell datasets were “condensed” by a first pass through Bookend Assemble in which no incomplete transcripts were discarded (FigS5A; see Supporting Notes: “Path Filtering”). Assembly was run again on the 96 condensed files, only retaining complete transcript models during the second pass. Bookend was also used to meta-assemble the 96 single-cell assemblies by StringTie2 and Scallop. Compared to standard assembly by StringTie2 or Scallop, all meta-assemblies produced substantially fewer non-matching transcripts (Fig5C). However, single-cell meta-assemblies surprisingly also recalled fewer RefSeq matches than standard assembly, with the exception of Bookend-to-Bookend and hybrid Bookend-to-Bookend meta-assemblies. PsiCLASS and TACO both showed somewhat higher precision than standard assembly, but at the expense of a severe drop in recall (Fig5D). PsiCLASS had the lowest recall of any method, but higher precision than StringTie2-to-TACO or Scallop-to-TACO meta-assembly. Bookend-to-Bookend meta-assemby considerably outperformed PsiCLASS in both recall (relative increase of 72%) and precision (relative increase of 253%). PsiCLASS produced an unusually large number of partial transcript fragments, likely due to the fact that scRNA-seq often has substantial 3′ bias that is not adequately accounted for (FigS5A-B). Notably, when TACO was applied to single-cell Bookend assemblies, it showed both a 23% relative reduction in recall and a 25% relative reduction in precision compared to standard Bookend assembly. In contrast, Bookend-to-Bookend meta-assembly increased recall by 22% and precision by 7% (+58% recall and +42% precision vs. Bookend-to-TACO). Across all three base assemblers, TACO reported fewer full reference matches than the standard assembly, while Bookend reported the same number or more full matches with a greater reduction in all non-matching classes than TACO (FigS5C). Of all combinations tested, both sensitivity and precision were highest at the intron chain and full transcript level in a Bookend-to-Bookend hybrid meta-assembly in which SLIC-CAGE and 3P-seq data were supplied alongside the single-cell condensed assemblies^55^ (Supplemental Table S5). We report this assembly as the “Bookend mESC” annotation (Supplemental Dataset 3-4). Requiring that both transcript ends are replicable across at least two different samples raised the transcript-level concordance with RefSeq to 54.1%, a relative increase of 271% over the most precise non-Bookend method (PsiCLASS), and a substantially higher agreement than even Gencode, an alternative mouse reference annotation that only shares 31.7% of its transcripts at assembled loci with RefSeq (FigS5C). While Gencode isoforms contain a broader set of alternative TSS and PAS than RefSeq, we noticed that they can be contained in low-confidence or fragmented transcript models, as in the gene Commd8 (Fig5E). By combining multiple unique advantages of end-guided assembly, Bookend could assemble more reference matches than any other strategy while maintaining a majority concordance with known annotations.

## DISCUSSION

Computational gene annotation pipelines have long struggled to produce a reliable picture of plant and animal transcriptomes at the isoform level^11,29,56^. Studying the details of gene regulation and isoform usage remains restricted to a small number of model organisms in which manually curated accurate transcript models are available. Even with specialized methods for sequencing RNA ends, connecting those ends to a gene model can be computationally challenging, especially for noncoding RNAs^35^. By generating accurate end-to-end transcript assemblies from a range of widely accessible sequencing methods, Bookend enables the automated annotation of promoter architecture, alternative polyadenylation and splicing dynamics in tissues in response to developmental, environmental and disease state cues.

Despite rapid advancements in scale and sensitivity of single-cell RNA sequencing, the accurate detection of transcript isoforms is still an outstanding challenge^48^. Full-length cDNA can be amplified from single cells with the Smart-seq family of “full-length sequencing” methods, including the recently developed Smart-seq3 that more efficiently captures 5’-labeled ends and gene body reads simultaneously^22,23^. Multiple approaches to apply long-read sequencing to single cells have been developed, but limits on throughput, error rate, and cost restricts their use^57–59^. Notably, large-scale Smart-seq2 experiments across multiple organisms have already been sequenced, including tens of thousands of cells from 20 mouse tissues and 24 human tissues by the Tabula Muris and Tabula Sapiens Consortia, respectively^60,61^. Through meta-assembly of full-length scRNA-seq data, Bookend enables the wholesale reannotation of genomes at single-cell resolution using existing and future datasets.

## METHODS

### PacBio Sequencing

Two PacBio Iso-seq libraries were generated each using 10 µg of total RNA from Arabidopsis inflorescences containing unopened floral buds. Total RNA was extracted with TRIzol following the method described in Schon et al. 2018^16^ to yield two biological replicates with an RNA integrity number (RIN) of 9.0 and 9.2, respectively. SMRTbell libraries were constructed by the Vienna BioCenter Core Facilities (VBCF) and sequenced on a Sequel SMRT Cell 1M.

### Published RNA sequencing data

Smart-seq2 datasets from 5ng Arabidopsis thaliana floral bud RNA and tissue-matched nanoPARE libraries from 10ug total RNA were downloaded from the NCBI Gene Expression Omnibus (GEO), series accession GSE112869. Single-cell RNA-seq of mouse embryonic stem cells and SIRVs from Natarajan et al. 2019 was downloaded from EMBL-EBI ArrayExpress, accession E-MTAB-7239. SLIC-CAGE samples from 100ng mESC total RNA were downloaded from ArrayExpress, accession E-MTAB-6519. One 3P-Seq library from 75ug mESC RNA was downloaded from GEO, sample accession GSM1268958.

### Short read data processing

Prior to alignment, reads were preprocessed with cutadapt^62^ to remove sequencing adapters. End labels were identified and trimmed using the utility *bookend label*, with settings tailored to each library. For Arabidopsis single-end Smart-seq2 reads, the arguments *--strand unstranded -S AAGCAGTGGTATCAACGCAGAGTACGGG –E AAGCAGTGGTATCAACGCAGAGTACTTTTTTTTTTTTTTTTTTTT+ --min_start 7 --min_end 9 - -minlen 18 --minqual 25 --qualmask 16 --mismatch_rate 0*.*06* were used. Paired-end mouse SMARTer reads used the same arguments except for *-S AAGCAGTGGTATCAACGCAGAGTACATGGG*. 5’ end reads from nanoPARE libraries were labeled with the arguments *--strand forward --minstart 20*. After end labeling, short reads were aligned using STAR^63^. Arabidopsis reads were aligned to the TAIR10 genome, and mouse reads were aligned to mm39 (GRCm39). Short reads in both species were aligned using an identical two-pass alignment strategy except for allowed intron lengths. First, reads were aligned with the command *STAR --runMode alignReads --alignEndsType EndToEnd --outFilterMatchNmin 20 --outFilterMismatchNmax 6 --outFilterMismatchNoverLmax* .*05 –outFilterIntronMotifs RemoveNoncanonicalUnannotated --alignSJoverhangMin 20 --alignSJDBoverhangMin 1 --outFilterMultimapNmax 2 --outSJfilterOverhangMin -1 15 20 20 --outSJfilterCountUniqueMin -1 2 3 3 --outSJfilterCountTotalMin -1 2 3 3*. Arabidopsis alignments used the additional arguments *--alignIntronMax 5000 --alignMatesGapMax 5100*, and mouse alignments instead used *--alignIntronMax 100000 --alignMatesGapMax 100100*. Splice junctions from all samples were aggregated across all samples for each species with *bookend sj-merge --new --min_reps 2* to retain only novel splice junctions that were detected in multiple samples. Second pass mapping was performed with the settings above, except the merged splice junction file was provided with *--sjdbFileChrStartEnd*, and the following arguments were modified: *--alignEndsType Local --outFilterMatchNminOverLread 0*.*9 --outFilterType BySJout --outFilterMultimapNmax 10 --outSAMtype BAM Unsorted --outSAMorder Paired --outSAMprimaryFlag AllBestScore --outSAMattributes NH HI AS nM NM MD jM jI XS*. Unsorted BAM files were converted to End-Labeled Read (ELR) files with the command *bookend elr --genome [genome*.*fa]* with library-specific settings. Arabidopsis Smart-seq2: *--start_seq ACGGG --end_seq RRRRRRRRRRRRRRRRRRRRRRRRRRRRRR --mismatch_rate* .*2;* Arabidopsis nanoPARE: --stranded *-s --start_seq ACGGG --mismatch_rate* .*2*; mouse SMARTer: *--start_seq ACATGGG --end_seq AAAAARRRRRRRRRRRRRRRRRRRRRRRRR --mismatch_rate* .*25*.

### Long read data processing

Raw Arabidopsis PacBio reads were converted to Circular Consensus Sequences using Iso-seq3 software with the command *ccs --min-passes 2 --min-rq* .*9*, and CCS reads were converted to full-length non-chimeric (FLNC) reads using *lima* and *isoseq3 refine --require-polya --min-rq -1 --min-polya-length 10*. FLNC reads were aligned to the Arabidopsis genome with the command *minimap2 -G 5000 -H -ax splice --MD -C 5 -u f -p 0*.*9 --junc-bed [TAIR10 transcript BED12]*. Aligned unsorted SAM files were converted to ELR with the command *bookend elr --stranded –s -e --start_seq ATGGG --genome [TAIR10*.*fa]*.

### Assembly

To make assembly setting maximally uniform across Bookend, StringTie2, Scallop, and Cufflinks, the following arguments were used. For Arabidopsis assemblies: *bookend --max_gap 50 --min_cov 2 --min_len 60 --min_proportion 0*.*02 --min_overhang 3 --cap_bonus 5 --cap_filter 0*.*02*; *stringtie -g 50 -c 2 -m 60 -f 0*.*02 -a 3 -M 1 -s 5; scallop --min_bundle_gap 50 --min_transcript_coverage 2 --min_transcript_length_base 60 --min_flank_length 3 --min_single_exon_coverage 5 --min_transcript_length increase 50; cufflinks -F 0*.*02 --overhang-tolerance 3 --min-frags-per-transfrag 10 -j 0*.*15 -A 0*.*06*. For mouse assemblies the same settings were used with the following exceptions: *--min_proportion* was set to 0.01, --min_len to 200, and *--require_cap* was enforced on mouse assemblies except when assembling spike-in transcripts, which do not possess caps. For meta-assembly, Bookend was run with the same settings as above for mouse. TACO was run with the arguments *--filter-min-expr 2 --filter-min-length 200 --isoform-frac 0*.*01*, and PsiCLASS was run with default settings

### Assembly algorithms

A brief overview of the end-guided assembly process implemented in Bookend is below. For a full breakdown of the algorithms used, see the “Bookend Algorithms” Supplemental Note. (*Generate Chunks*) First, reads are streamed in from an ELR file in sorted order and separated into overlapping chunks. (*Tag Clustering*) In each chunk, Start Tags and End Tags are clustered on each strand by grouping tags by genomic position and assigning each position a signal score of counts ×proportion of total coverage. A signal threshold is set and positions below the threshold are discarded. Remaining positions are grouped within a user-specified distance to yield Start and End clusters on each strand. (*Calculate Membership Matrix*) Start/End clusters are added to a catalog of boundaries, which include splice donor/acceptor sites that are also filtered by a threshold of total overlapping coverage. Adjacent boundary pairs define a “frag”, and each read is assigned a Membership array that describes whether the read overlaps or excludes each frag. Redundant membership arrays are combined, and the unique set of elements is stored as the Membership Matrix. (*Calculate Overlap Matrix*) A matrix describing the relationship between each element pair *a* and *b* is generated by asking (from left to right in genomic coordinates): can *a* extend into *b*? Can *b* extend into *a*? Each comparison returns a pair of Overlaps, *Oab* and *Oba*, respectively: 1 = extends, -1 = excludes, 2 = is contained by, 0 = does not overlap. The values - 1 and 0 are symmetric, but 1 and 2 are directed relationships that can be used as edges in a directed graph. (*Collapse Linear Chains*) It is possible to identify and collapse non-branching sets of elements (“linear chains”) prior to assembly. Two graphs are constructed with elements as nodes: a directed graph with extensions as edges, and an undirected graph with exclusions as edges. A depth-first search is conducted by visiting each element in increasing order of information content (number of non-zero memberships). During a visit, the element’s edges are traversed recursively to record all traversed nodes’ exclusions. An element with no edges is assigned to a new chain. Otherwise, when an element’s edges are all traversed, the element is compared against its outgroup, the set of all elements reached. If all outgroup elements belong to one chain and the element and outgroup have the same set of exclusions, then the element is added to the same chain. If the element’s outgroup is assigned to multiple chains, the element begins a new chain. After completion of the search, each chain is combined to form a single reduced element. (*Generate Overlap Graph*) From the set of reduced elements a second directed graph is constructed with a global source (Start+/End-) and sink (Start-/End+), where each node records the element weight (sequenced bases / genomic length), outgroup (extends to), ingroup (extends from), containments and exclusions. (*Resolve Containment*) All elements contained by one or more longer elements have their weight redistributed proportionally to their container as long as not all containers exclude any single node the element doesn’t already exclude. (*Greedy Paths*) All elements begin unassigned. Starting with the heaviest unassigned element, choose an extension (ingroup/outgroup pair) that maximizes a score that equally combines the following: maximal weight of the extension, maximal similarity of weight distribution across samples between element and extension, minimal coverage variance across covered frags, and does not cause the source or sink to become unreachable. The highest-scoring extension is iteratively added to a path until both source and sink are reached. Paths are generated in this manner until the total weight of unassigned elements falls below a given signal threshold.

## Supporting information

Suppemental Materials

## Contributions

M.A.S. and M.D.N. conceived the project; M.A.S. developed the methodology; M.A.S and S.L. performed the experiments; M.A.S. and F.H. analyzed data; M.A.S. prepared figures; M.A.S wrote the article; M.A.S. and M.D.N. edited the article; M.D.N. acquired funding and supervised the project.

## Acknowledgements

We thank the Next Generation Sequencing Facility at Vienna BioCenter Core Facilities GmbH (VBCF) for their outstanding services and technical support.

## Data access

Bookend software is available on the Python Package Index and can be installed with the command *pip install bookend-rna*. Source code is available as a repository on GitHub at https://github.com/Gregor-Mendel-Institute/bookend. All sequencing data generated in this study have been submitted to the National Center for Biotechnology Information Gene Expression Omnibus (NCBI GEO, https://www.ncbi.nlm.nih.gov/geo/) under accession number GSE189482.

