## Supplementary material for "Precise Transcript Reconstruction with End-Guided Assembly": Suppemental Materials

### SUPPLEMENTAL MATERIAL

#### Precise Transcript Reconstruction from Single Cells via End-Guided Assembly

This document serves as an Appendix to [Schon et al. 2021], and it contains descriptions of algorithms used by the end-guided transcript assembler Bookend. Source code can be found at https://github.com/Gregor-Mendel-Institute/bookend.

The package can be installed from the Python Package Index on any system with Python 3.6+ using the command `pip install bookend-rna`.

Supplemental Datasets are available as a separate Excel spreadsheet:

| | |
|---|---|
| Data S1. | Bookend Floral Bud- hybrid assembly Arabidopsis stage 12 inflorescence |
| Data S2. | Classification of Bookend Floral Bud transcripts against TAIR10 and Araport11 |
| Data S3. | Bookend mESC- hybrid assembly of mouse embryonic stem cells |
| Data S4. | Classification of Bookend mESC transcripts against RefSeq and Gencode |

### SUPPLEMENTAL FIGURES

#### Figure S1. The Bookend workflow

**A** Workflow diagram showing the computational steps to generate an end-guided assembly from raw FASTQ RNA-seq files with Bookend. **B** Schematic of the Membership Matrix produced from a hypothetical collection of reads from various sequencing methods; blue- frag is included, pink- frag is excluded. Elements that include or exclude every frag are considered complete, e.g. the first and last long reads.

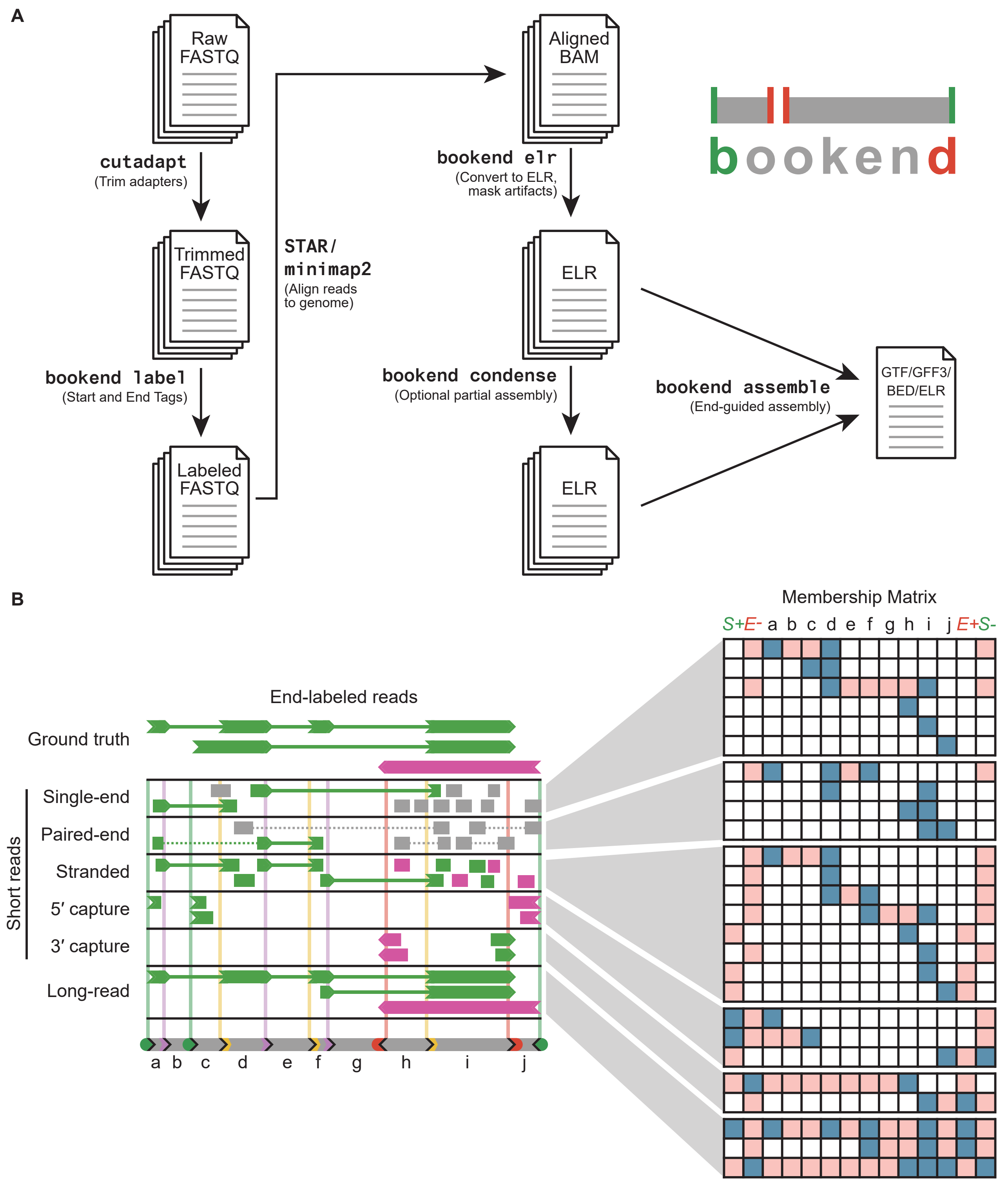

#### Figure S2. Nucleotide-level precision of Arabidopsis assembly 5′ and 3′ ends

**A** Percent of the unique set of 5′ ends from different assemblies of floral bud RNA, or from reference annotations (TAIR10, Araport11) overlapping assembled loci, that fall on or adjacent to the peak position of a CAGE cluster from Thieffry et al. 2020. **B** Validation as in **A** for the set of unique 3′ ends against Direct RNA-Seq (DRS) data from Sherstnev et al. 2012. **C** (Top) Nucleotide fold enrichment over background frequencies in a $\pm$50nt window around the peak positions of all CAGE clusters; (bottom left) nucleotide enrichment around assembly 5′ ends $\leq$ 100nt from a TAIR10 TSS; (bottom right) enrichment around all other assembly 5′ ends. **D** Enrichments displayed as in **C** for DRS clusters and assembly 3′ ends. **E** Estimated read coverage distribution for assembled transcripts grouped by their classification against TAIR10. Center line- median coverage. Outliers are not shown.

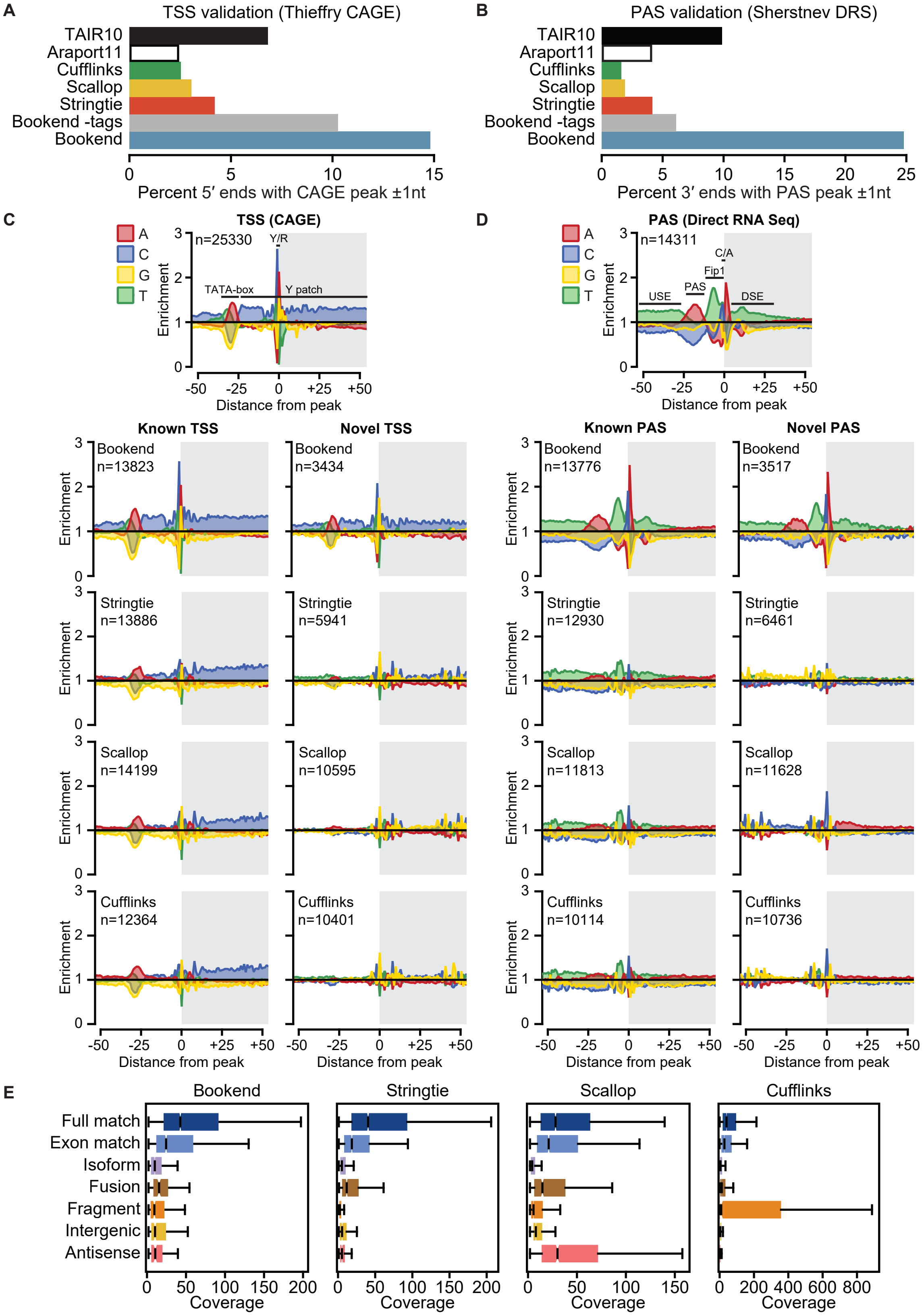

#### Figure S3. Artifacts in long-read data

IGV browser image of the Arabidopsis *MIA* locus. From top to bottom: PacBio FLNC reads, colored by 3′ label type; Bars demarcating genomic regions of 10 or more consecutive A's (10 or more T's on the reverse strand); Direct RNA Seq (DRS) 3′ end abundance; nanoPARE 5′ end capped and noncapped read abundance; Smart-seq2 coverage depth; TAIR10 reference; Assemblies colored by class vs. TAIR10. **B** (Upper panel) Nucleotide frequency enrichment in a ±50bp window around transcription start sites (TSS) identified by CAGE (Thieffry et al. 2020). (Lower panel) Nucleotide enrichment around 5′ ends of transcripts constructed from PacBio reads by Iso-seq3 (top), Stringtie2 (middle), and Bookend (bottom) at sites overlapping a TAIR10 TSS (left), novel TSS at a known gene (middle), and novel antisense or intergenic loci (right). **C** Precision/recall plot of assemblers on Smart-seq2 (short reads) and/or Pacbio (long reads).

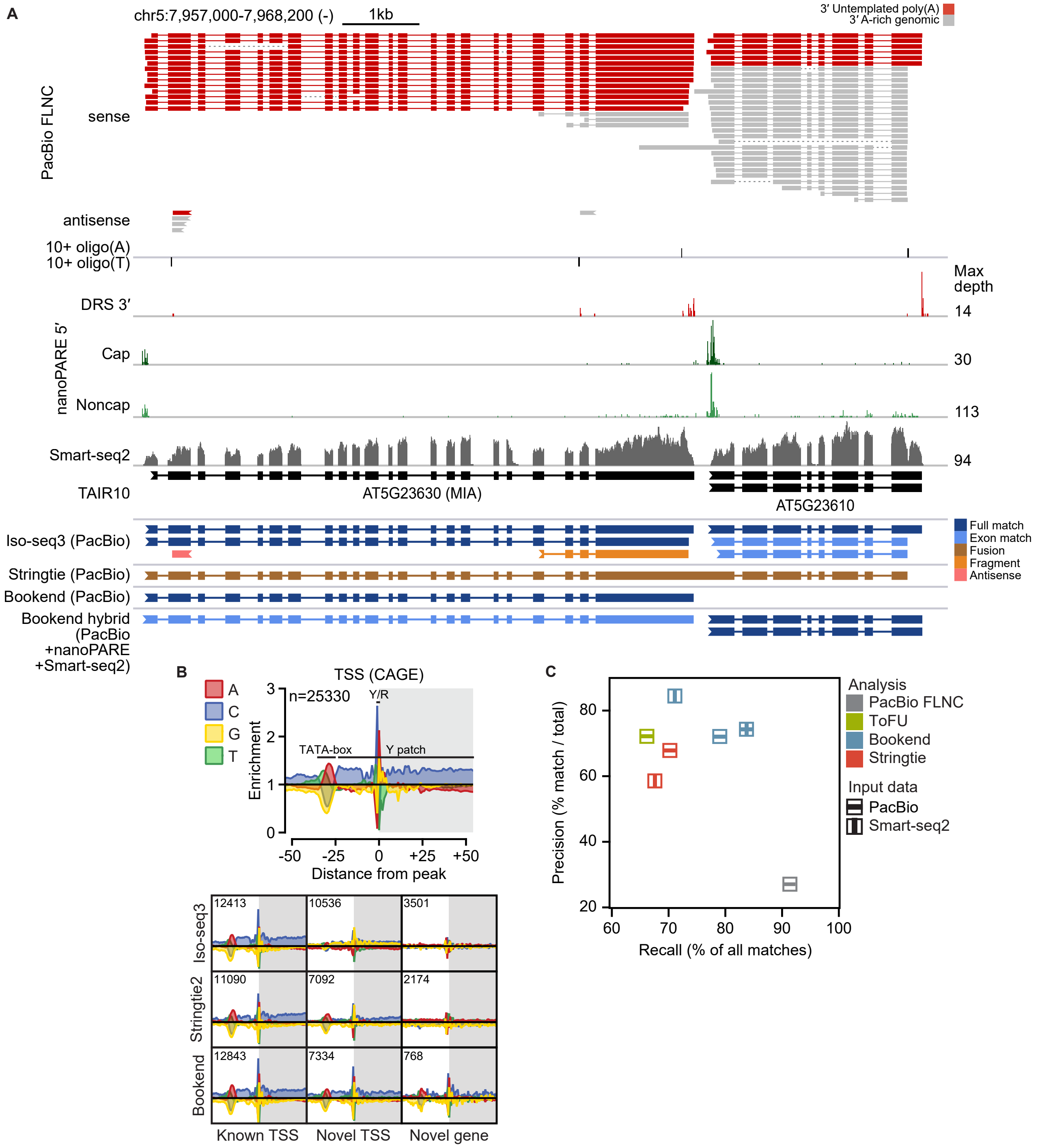

#### Figure S4. Single mESC assembly details

**A** IGV browser image of the synthetic SIRV6 locus. (Top) Mix E2 spike-in concentrations for 18 distinct RNA molecules. (Bottom) Assembled isoforms of pooled data from Natarajan et al. 2019 of 96 SMARTer libraries of single mouse embryonic stem cells with SIRV spike-ins. Transcripts are colored by the concentration of their matching SIRV transcript, if it exists. **B** Precision/recall plots for assemblies of 96 individual mESCs whose RNA was split to generate three RNA-seq libraries from two different sequencing protocols. Black points incorporate mESC SLIC-CAGE data from Cvetesic et al. 2018 and mESC 3P-Seq data from Nam et al. 2013 in a hybrid assembly.

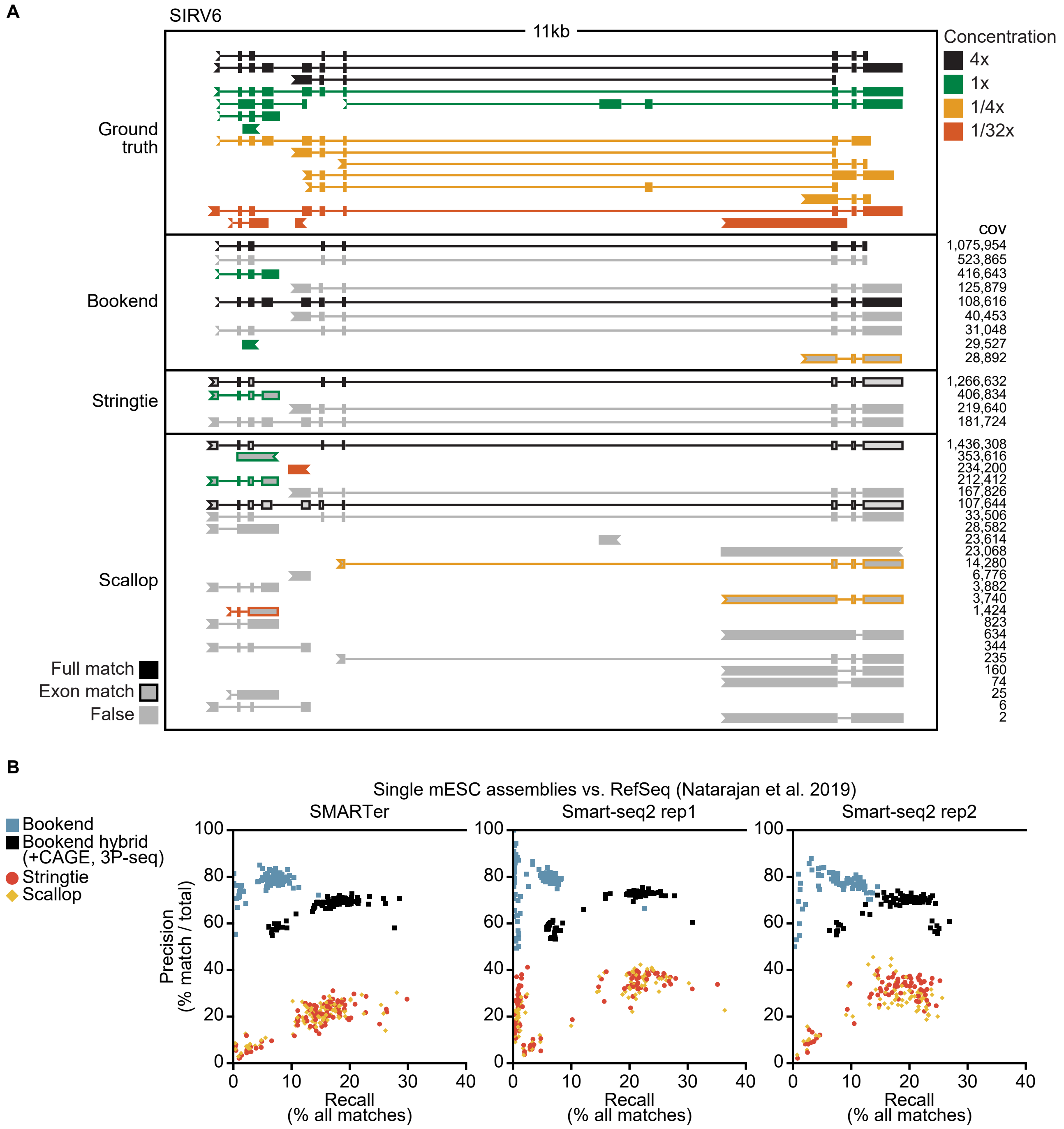

#### Figure S5. Meta-assembly details

**A** IGV browser image of the mouse Gga1 gene. Tracks from top to bottom: read abundance of mESC SLIC-CAGE and 3P-seq; Bookend partial assembly of SMARTer data from 96 mESC cells; meta-assembly by Bookend, PsiClass, and TACO. **B** Classification of all assembled transcripts with and without meta-assembly. **C** Change in transcript abundance by class after meta-assembly by TACO or Bookend. **D** Euler diagram of the number of shared exon chains between Bookend mESC, RefSeq and Gencode annotations at loci assembled by Bookend.

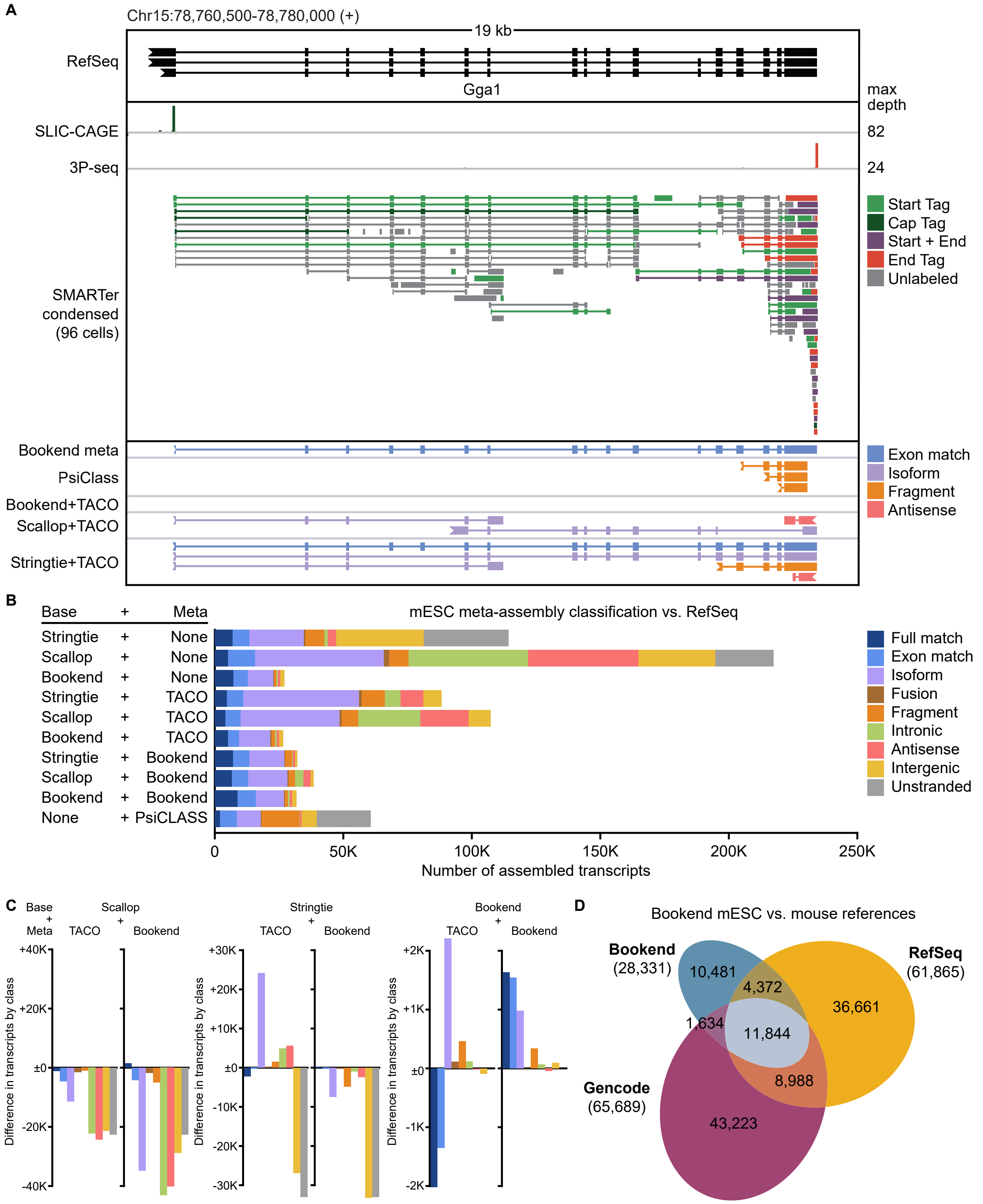

### SUPPLEMENTAL TABLES

#### Table S1. Floral bud Smart-seq2 end-labeled read mapping statistics

End-labeled reads identified in three Smart-seq2 replicates from 5ng of Arabidopsis floral bud total RNA, sequenced single-end 50bp mode on an Illumina Hi-Seq 2500.

LABEL

| Read type | Rep 1 | Rep 2 | Rep 3 | Merged |
|---|---|---|---|---|
| Raw reads | 26,714,548 | 24,104,566 | 24,626,064 | 75,445,178 |
| 5′ Label | 743,292 | 948,127 | 657,093 | 2,348,512 |
| 3′ Label | 194,038 | 389,972 | 211,447 | 795,457 |

ALIGN

| Read type | Rep 1 | Rep 2 | Rep 3 | Merged |
|---|---|---|---|---|
| Unique mappers | 20,218,104 (76%) | 16,552,901 (69%) | 18,655,520 (76%) | 55,426,525 (73%) |
| Start Tag | 446,529 (60%) | 523,009 (55%) | 416,070 (63%) | 1,385,608 (59%) |
| End Tag | 126,315 (65%) | 252,673 (65%) | 149,980 (71%) | 528,968 (66%) |
| Multi-mappers | 5,266,353 (20%) | 5,609,624 (23%) | 4,645,067 (19%) | 15,521,044 (21%) |
| Start Tag | 213,413 (29%) | 273,929 (29%) | 165,207 (25%) | 652,549 (28%) |
| End Tag | 31,979 (16%) | 53,612 (14%) | 23,970 (11%) | 109,561 (14%) |

FILTER

| Read type | Rep 1 | Rep 2 | Rep 3 | Merged |
|---|---|---|---|---|
| 5′ TSO artifacts | 71,921 (11%) | 108,081 (14%) | 74,085 (13%) | 254,087 (12%) |
| 5′ uuG Caps | 51,748 (8%) | 48,398 (6%) | 39,601 (7%) | 139,747 (7%) |
| 3′ poly(A) artifacts | 52,047 (33%) | 91,573 (30%) | 50,166 (29%) | 193,786 (30%) |

FINAL

| Read type | Rep 1 | Rep 2 | Rep 3 | Merged |
|---|---|---|---|---|
| Unlabeled | 24,666,221 | 21,059,302 | 22,545,360 | 68,270,883 |
| Start Tags | 536,273 | 640,459 | 467,591 | 1,644,323 |
| Cap Tags | 51,748 | 48,398 | 39,601 | 139,747 |
| End Tags | 126,315 | 214,712 | 123,784 | 464,811 |

#### Table S2. Long-read validation of floral bud assemblies by class

Breakdown of assembled transcripts by their classification against the closest TAIR10 reference transcript, and the number of transcripts in each class that have at least one full match (exon chain and ±100 ends) in PacBio long-read sequencing data from 10ug of *Arabidopsis thaliana* stage 12 floral bud total RNA.

CLASSIFICATION

| Class | Bookend | Bookend -tags | Stringtie | Scallop | Cufflinks |
|---|---|---|---|---|---|
| Full match | **12019** | 7809 | 10674 | 9068 | 5480 |
| Exon match | 4139 | 5851 | 4579 | **6717** | 5571 |
| Isoform | 1915 | 2409 | 5337 | 8317 | **9728** |
| Fusion | 185 | 1101 | 1049 | 1439 | **3737** |
| Fragment | 362 | 1325 | 1813 | 3431 | **3955** |
| Intergenic | 399 | 279 | 530 | 1113 | **1160** |
| Antisense | 112 | 70 | 145 | **3049** | 341 |
| Intronic | 6 | 3 | 12 | **51** | 13 |
| Ambiguous | 0 | 819 | 1910 | **50185** | 6025 |
| Known | **16158** | 13660 | 15253 | 15785 | 11051 |
| Novel | 2979 | 5187 | 8886 | 17400 | **18934** |
| Total | 19137 | 18847 | 24139 | **33185** | 29985 |

VALIDATION

| Class | Bookend | Bookend (-tags) | Stringtie | Scallop | Cufflinks |
|---|---|---|---|---|---|
| Full match | **11239 (94%)** | 7070 (91%) | 9671 (91%) | 7925 (87%) | 4619 (84%) |
| Exon match | **3033 (73%)** | 2450 (42%) | 2124 (46%) | 1961 (29%) | 1252 (23%) |
| Isoform | **903 (47%)** | 660 (27%) | 1216 (23%) | 1630 (20%) | 693 (7%) |
| Fusion | **56 (30%)** | 42 (4%) | 60 (6%) | 87 (6%) | 36 (1%) |
| Fragment | **116 (32%)** | 63 (8%) | 80 (4%) | 83 (2%) | 65 (2%) |
| Intergenic | **147 (37%)** | 21 (8%) | 46 (9%) | 68 (6%) | 20 (2%) |
| Antisense | **38 (34%)** | 5 (7%) | 7 (5%) | 150 (5%) | 9 (3%) |
| Intronic | **1 (17%)** | 0 (0%) | 0 (0%) | 2 (4%) | 0 (0%) |
| Ambiguous | N/A | N/A | N/A | N/A | N/A |
| Known | **14272 (88%)** | 9520 (70%) | 11795 (77%) | 9886 (63%) | 5871 (53%) |
| Novel | **1261 (42%)** | 791 (15%) | 1409 (16%) | 2020 (12%) | 823 (4%) |
| Total | **15533 (81%)** | 10311 (55%) | 13204 (55%) | 11906 (36%) | 6694 (22%) |

#### Table S3. Floral bud hybrid assembly details

Classification against TAIR10 of transcripts assembled from a set of complementary datasets from floral bud RNA. Short reads- Smart-seq2; Start reads- nanoPARE; Long reads- PacBio.

LONG READS ONLY

| Class | PacBio FLNC | (unique) | Iso-seq3 | (unique) | ToFU | Stringtie | Bookend |
|---|---|---|---|---|---|---|---|
| Full match | 283936 | (14136) | 21873 | (11927) | 12596 | 10892 | **13357** |
| Exon match | 151425 | (7221) | 12525 | (3364) | **5217** | 4986 | 4946 |
| Isoform | 39863 | (23055) | 4260 | (3551) | 3177 | 4089 | **5546** |
| Fusion | 3513 | (803) | 341 | (131) | 163 | **641** | 294 |
| Fragment | 47986 | (19459) | 7150 | **(4824)** | 638 | 539 | 400 |
| Intergenic | 35620 | (9571) | 3336 | **(2117)** | 2065 | 1760 | 394 |
| Antisense | 7682 | (4465) | 929 | **(817)** | 803 | 481 | 438 |
| Intronic | 544 | (155) | 60 | **(30)** | **30** | 11 | 9 |
| Known | 435361 | (21357) | 34398 | (15291) | 17813 | 15878 | **18303** |
| Novel | 135208 | (57508) | 16076 | **(11470)** | 6876 | 7521 | 7081 |
| Total | 570569 | (78865) | 50474 | **(26761)** | 22754 | 23399 | 25384 |
| (% Known) | 76.3% | (27.1%) | 68.1% | (57.1%) | 69.8 | 67.9% | **72.1%** |

BOOKEND HYBRID ASSEMBLY

| Class | Long+Short | Long+Start | Short+Start | Long+Short+Start | (require cap) |
|---|---|---|---|---|---|
| Full match | **14166** | 12915 | 12183 | 14098 | (14132) |
| Exon match | 5452 | 4434 | 4435 | **5559** | (5161) |
| Isoform | 5050 | 4977 | 2199 | **5387** | (5268) |
| Fusion | **348** | 205 | 141 | 276 | (258) |
| Fragment | 321 | 399 | **536** | 434 | (258) |
| Intergenic | 579 | 395 | 411 | **639** | (414) |
| Antisense | **476** | 384 | 122 | 443 | (364) |
| Intronic | 9 | 9 | 8 | **10** | (6) |
| Known | 19618 | 17349 | 16618 | **19657** | (19293) |
| Novel | 6783 | 6369 | 3417 | **7189** | (6568) |
| Total | 26401 | 23718 | 20035 | **26846** | (25861) |
| (% Known) | 74.3% | 73.1% | 82.9% | 73.2% | (74.6%) |

#### Table S4. End-labeling and alignment of single mESCs

Mapping statistics for 96 single mESC SMARTer libraries from Natarajan et al. 2019.

| Cell | Unlabeled | 5′ label | 3′ label | Aligned | Start Tag | Cap Tag | (%) | End Tag | (%) |
|---|---|---|---|---|---|---|---|---|---|
| 1 | 9608668 | 439193 | 100121 | 8934461 | 258330 | 79340 | 0.9% | 37906 | 0.4% |
| 2 | 9831074 | 618059 | 118804 | 9607068 | 365454 | 96924 | 1.0% | 54055 | 0.6% |
| 3 | 12146979 | 641146 | 127258 | 11425200 | 386469 | 105849 | 0.9% | 51691 | 0.5% |
| 4 | 6335181 | 298745 | 79348 | 5750133 | 178094 | 51310 | 0.9% | 31671 | 0.6% |
| 5 | 7514264 | 427582 | 90844 | 7177000 | 251555 | 73783 | 1.0% | 40753 | 0.6% |
| 6 | 6192857 | 424987 | 71214 | 5568044 | 238054 | 64005 | 1.1% | 29469 | 0.5% |
| 7 | 9413990 | 557275 | 105944 | 8676900 | 308285 | 97282 | 1.1% | 41137 | 0.5% |
| 8 | 7146811 | 478394 | 109538 | 6626337 | 303099 | 55831 | 0.8% | 41087 | 0.6% |
| 9 | 12209527 | 656470 | 119208 | 11653781 | 364251 | 126662 | 1.1% | 51548 | 0.4% |
| 10 | 4943634 | 249114 | 57765 | 4150020 | 133363 | 44176 | 1.1% | 22076 | 0.5% |
| 11 | 10289384 | 579612 | 156501 | 9540995 | 344615 | 107138 | 1.1% | 67970 | 0.7% |
| 12 | 6296290 | 354539 | 36954 | 5744670 | 207542 | 57459 | 1.0% | 15386 | 0.3% |
| 13 | 11062529 | 608742 | 107064 | 10588553 | 402291 | 90084 | 0.9% | 38529 | 0.4% |
| 14 | 5859896 | 372904 | 57864 | 5597543 | 221374 | 58681 | 1.0% | 25575 | 0.5% |
| 15 | 7438431 | 377316 | 85287 | 6600211 | 216701 | 60159 | 0.9% | 31166 | 0.5% |
| 16 | 9318662 | 489053 | 77267 | 8804817 | 287566 | 85378 | 1.0% | 29135 | 0.3% |
| 17 | 9654938 | 591123 | 111696 | 8821047 | 345010 | 95892 | 1.1% | 46811 | 0.5% |
| 18 | 4176594 | 309959 | 77647 | 3386566 | 196545 | 14120 | 0.4% | 21449 | 0.6% |
| 19 | 10166487 | 646012 | 87377 | 9584997 | 400128 | 99395 | 1.0% | 34685 | 0.4% |
| 20 | 4513269 | 374249 | 70382 | 3592280 | 232724 | 13498 | 0.4% | 17983 | 0.5% |
| 21 | 5911031 | 367655 | 77492 | 5715353 | 230948 | 68676 | 1.2% | 34725 | 0.6% |
| 22 | 8533714 | 500707 | 100164 | 8057641 | 293859 | 101787 | 1.3% | 44715 | 0.6% |
| 23 | 5507278 | 446779 | 71288 | 4598916 | 301960 | 17407 | 0.4% | 19768 | 0.4% |
| 24 | 7007715 | 436446 | 81182 | 6404774 | 243859 | 75132 | 1.2% | 33581 | 0.5% |
| 25 | 3995373 | 323171 | 74361 | 3632465 | 237703 | 10268 | 0.3% | 20365 | 0.6% |
| 26 | 5412175 | 277064 | 69361 | 5014494 | 165316 | 45054 | 0.9% | 25527 | 0.5% |
| 27 | 4498372 | 268910 | 56498 | 4148402 | 168253 | 40980 | 1.0% | 23122 | 0.6% |
| 28 | 29923526 | 1235689 | 302270 | 29149150 | 791759 | 206991 | 0.7% | 117971 | 0.4% |
| 29 | 10165433 | 565658 | 123935 | 9849721 | 358781 | 93532 | 0.9% | 48662 | 0.5% |
| 30 | 6088334 | 309792 | 72381 | 5791867 | 198551 | 44964 | 0.8% | 24476 | 0.4% |
| 31 | 7560915 | 419014 | 82924 | 6738642 | 265942 | 45899 | 0.7% | 25400 | 0.4% |
| 32 | 5485800 | 345511 | 78723 | 4856333 | 199379 | 50828 | 1.0% | 29350 | 0.6% |
| 33 | 5373713 | 373940 | 66041 | 4823649 | 224204 | 52800 | 1.1% | 29194 | 0.6% |
| 34 | 6485920 | 426953 | 71948 | 5608916 | 225477 | 65596 | 1.2% | 29468 | 0.5% |
| 35 | 6632255 | 360820 | 74635 | 5934034 | 219050 | 49897 | 0.8% | 26091 | 0.4% |
| 36 | 5977435 | 506299 | 90559 | 5824146 | 368469 | 27634 | 0.5% | 32330 | 0.6% |
| 37 | 2705467 | 189226 | 58195 | 1931832 | 118339 | 4561 | 0.2% | 28556 | 1.5% |
| 38 | 6035540 | 371889 | 79129 | 5753194 | 233286 | 64046 | 1.1% | 36886 | 0.6% |
| 39 | 4425629 | 292098 | 59420 | 4081161 | 168990 | 48537 | 1.2% | 26558 | 0.7% |
| 40 | 6388799 | 427916 | 66878 | 6408595 | 301594 | 60211 | 0.9% | 32197 | 0.5% |
| 41 | 7985364 | 460244 | 115172 | 7245203 | 269110 | 82244 | 1.1% | 48552 | 0.7% |
| 42 | 7125365 | 342965 | 73899 | 6819032 | 226665 | 48831 | 0.7% | 23736 | 0.3% |
| 43 | 3268434 | 287751 | 64027 | 1932186 | 146951 | 5797 | 0.3% | 25172 | 1.3% |
| 44 | 6776069 | 348272 | 70412 | 5796199 | 198263 | 51087 | 0.9% | 26426 | 0.5% |
| 45 | 6267559 | 334111 | 94890 | 5700986 | 202717 | 57249 | 1.0% | 38813 | 0.7% |
| 46 | 7858871 | 514789 | 85073 | 7257444 | 324611 | 81400 | 1.1% | 35532 | 0.5% |
| 47 | 6456318 | 295115 | 47795 | 5819580 | 171799 | 50144 | 0.9% | 17653 | 0.3% |
| 48 | 6404736 | 411525 | 74035 | 5627612 | 232133 | 63004 | 1.1% | 29733 | 0.5% |

Table S4, continued.

| Cell | Unlabeled | 5′ label | 3′ label | Aligned | Start Tag | Cap Tag | (%) | End Tag | (%) |
|---|---|---|---|---|---|---|---|---|---|
| 49 | 3389101 | 340785 | 115033 | 3422875 | 284973 | 8238 | 0.2% | 33683 | 1.0% |
| 50 | 5667716 | 363316 | 70312 | 4977135 | 222168 | 45158 | 0.9% | 24698 | 0.5% |
| 51 | 2690373 | 227903 | 58683 | 1946712 | 141653 | 5730 | 0.3% | 29509 | 1.5% |
| 52 | 6928921 | 362190 | 107121 | 6545706 | 225017 | 60246 | 0.9% | 40119 | 0.6% |
| 53 | 9835754 | 726933 | 136436 | 9463908 | 428107 | 107183 | 1.1% | 62678 | 0.7% |
| 54 | 3011944 | 270978 | 65486 | 2543453 | 185334 | 13107 | 0.5% | 29921 | 1.2% |
| 55 | 5142534 | 484427 | 92112 | 5059348 | 385759 | 18370 | 0.4% | 36733 | 0.7% |
| 56 | 7140952 | 395151 | 86458 | 6594524 | 251851 | 60822 | 0.9% | 34039 | 0.5% |
| 57 | 6466893 | 413749 | 63792 | 5598598 | 217828 | 69633 | 1.2% | 29085 | 0.5% |
| 58 | 3300888 | 266911 | 71045 | 2541725 | 166731 | 8887 | 0.3% | 18473 | 0.7% |
| 59 | 5043344 | 309051 | 46246 | 4926209 | 192103 | 48126 | 1.0% | 20633 | 0.4% |
| 60 | 6986624 | 481048 | 80193 | 6427797 | 279353 | 77828 | 1.2% | 36246 | 0.6% |
| 61 | 3351952 | 234887 | 76558 | 3319998 | 176381 | 26927 | 0.8% | 41700 | 1.3% |
| 62 | 3396729 | 339973 | 85217 | 3274103 | 264194 | 12512 | 0.4% | 33795 | 1.0% |
| 63 | 6461789 | 421328 | 66538 | 5632766 | 236839 | 57459 | 1.0% | 26779 | 0.5% |
| 64 | 6404975 | 422427 | 85385 | 5785467 | 270948 | 41527 | 0.7% | 26209 | 0.5% |
| 65 | 6557014 | 342169 | 117742 | 5717291 | 193178 | 52509 | 0.9% | 42129 | 0.7% |
| 66 | 3335945 | 307264 | 73995 | 2564465 | 184833 | 11539 | 0.4% | 22667 | 0.9% |
| 67 | 6375049 | 322527 | 59493 | 5944082 | 199964 | 47648 | 0.8% | 22887 | 0.4% |
| 68 | 8177038 | 404930 | 93646 | 7564092 | 252086 | 70170 | 0.9% | 34101 | 0.5% |
| 69 | 5485331 | 301692 | 84083 | 4864549 | 167140 | 47594 | 1.0% | 32817 | 0.7% |
| 70 | 14155483 | 792368 | 132139 | 13194440 | 467058 | 123415 | 0.9% | 55436 | 0.4% |
| 71 | 7022871 | 439562 | 91457 | 6477060 | 283655 | 61701 | 1.0% | 33721 | 0.5% |
| 72 | 7114718 | 466219 | 72834 | 6382720 | 262582 | 74644 | 1.2% | 31258 | 0.5% |
| 73 | 7494258 | 359929 | 110643 | 7266395 | 224032 | 63724 | 0.9% | 45323 | 0.6% |
| 74 | 9200518 | 599103 | 116324 | 8846834 | 377761 | 89071 | 1.0% | 49789 | 0.6% |
| 75 | 4351846 | 413743 | 90042 | 4214629 | 315500 | 19554 | 0.5% | 33950 | 0.8% |
| 76 | 4062440 | 346688 | 83752 | 3356938 | 233841 | 12302 | 0.4% | 24880 | 0.7% |
| 77 | 6264611 | 353118 | 100568 | 5758030 | 210097 | 62049 | 1.1% | 43787 | 0.8% |
| 78 | 6425634 | 332547 | 63487 | 5703753 | 202498 | 45756 | 0.8% | 23191 | 0.4% |
| 79 | 7234869 | 381668 | 113175 | 6540410 | 221080 | 65323 | 1.0% | 48511 | 0.7% |
| 80 | 10168955 | 356816 | 144538 | 8923837 | 176870 | 53756 | 0.6% | 51863 | 0.6% |
| 81 | 6613647 | 359383 | 55306 | 6380549 | 217473 | 63718 | 1.0% | 26495 | 0.4% |
| 82 | 3280612 | 308671 | 87547 | 2792536 | 216501 | 8298 | 0.3% | 23508 | 0.8% |
| 83 | 6292413 | 380969 | 59590 | 5645459 | 212510 | 67444 | 1.2% | 25334 | 0.4% |
| 84 | 5619685 | 508203 | 75195 | 5003815 | 349041 | 31826 | 0.6% | 25397 | 0.5% |
| 85 | 9277709 | 604069 | 92078 | 9040233 | 420256 | 79640 | 0.9% | 41817 | 0.5% |
| 86 | 6115076 | 342837 | 118423 | 5611657 | 200182 | 62551 | 1.1% | 48331 | 0.9% |
| 87 | 8466897 | 479130 | 137519 | 8090541 | 289579 | 83847 | 1.0% | 56166 | 0.7% |
| 88 | 5910776 | 308345 | 101811 | 5749104 | 198101 | 53210 | 0.9% | 41257 | 0.7% |
| 89 | 8795051 | 480325 | 131394 | 7941499 | 273086 | 88094 | 1.1% | 49065 | 0.6% |
| 90 | 6935880 | 397900 | 130562 | 6470810 | 230171 | 70897 | 1.1% | 54555 | 0.8% |
| 91 | 8114433 | 474650 | 129503 | 7215724 | 267064 | 79081 | 1.1% | 45991 | 0.6% |
| 92 | 3884626 | 214058 | 72777 | 3214197 | 115207 | 34069 | 1.1% | 27498 | 0.9% |
| 93 | 8932179 | 552463 | 99937 | 8764917 | 340984 | 98729 | 1.1% | 44467 | 0.5% |
| 94 | 8641039 | 533914 | 100584 | 7795662 | 283006 | 100413 | 1.3% | 42540 | 0.5% |
| 95 | 6281335 | 391868 | 78347 | 5496105 | 206090 | 62725 | 1.1% | 29948 | 0.5% |
| 96 | 5000892 | 288069 | 78217 | 4875119 | 174688 | 53544 | 1.1% | 34999 | 0.7% |

#### Table S5. GffCompare performance statistics for mESC meta-assemblies

Comparison of mESC assemblies with and without meta-assembly vs. overlapping RefSeq transcripts. Command-line argument: `gffcompare -R --strict-match --no-merge -e 100 -s GRCm39.fasta -r [RefSeq] [Assembly]`

SENSITIVITY

| Base assembler | Meta-assembler | Nucleotide | Exon | Intron | Intron chain | Transcript |
|---|---|---|---|---|---|---|
| PsiCLASS | PsiCLASS | 41.6 | 44.0 | 49.0 | 13.0 | 13.1 |
| Stringtie | None | 50.3 | 55.1 | 59.5 | 15.4 | 15.4 |
| Stringtie | TACO | 43.3 | 53.7 | 58.4 | 13.3 | 13.1 |
| Stringtie | Bookend | 49.6 | 63.8 | 66.2 | 21.3 | 21.1 |
| Scallop | None | **58.9** | 54.1 | 60.5 | 17.0 | 16.9 |
| Scallop | TACO | 45.7 | 53.4 | 57.6 | 11.6 | 11.5 |
| Scallop | Bookend | 49.5 | 62.1 | 64.5 | 20.8 | 20.7 |
| Bookend | None | 58.1 | 67.4 | 70.0 | 20.0 | 20.1 |
| Bookend | TACO | 46.3 | 57.9 | 59.4 | 18.0 | 18.0 |
| Bookend | Bookend | 56.6 | **67.8** | **70.4** | **24.7** | **24.7** |
| Bookend+ | None | 57.4 | 67.1 | 69.7 | 19.8 | 19.9 |
| Bookend+ | Bookend | 56.7 | 68.7 | 71.5 | 25.0 | 25.0 |
| Bookend+ | Bookend (stringent) | 57.9 | 70.2 | 73.0 | 25.5 | 25.5 |

+ Hybrid assembly with mESC CAGE and 3P-seq data

PRECISION

| Base assembler | Meta-assembler | Nucleotide | Exon | Intron | Intron chain | Transcript |
|---|---|---|---|---|---|---|
| PsiCLASS | PsiCLASS | 36.7 | 61.0 | **95.8** | 28.0 | 14.6 |
| Stringtie | None | 32.3 | 52.0 | 86.2 | 29.6 | 12.1 |
| Stringtie | TACO | 38.8 | 53.0 | 78.8 | 15.9 | 12.1 |
| Stringtie | Bookend | **77.3** | 79.9 | 92.3 | 38.6 | 38.4 |
| Scallop | None | 16.1 | 35.4 | 82.2 | 21.5 | 6.8 |
| Scallop | TACO | 28.6 | 48.0 | 83.1 | 18.2 | 8.6 |
| Scallop | Bookend | 59.6 | 74.6 | 92.5 | 38.1 | 30.5 |
| Bookend | None | 56.8 | 84.1 | 95.0 | 50.8 | 44.7 |
| Bookend | TACO | 67.2 | 79.3 | 90.4 | 40.8 | 35.4 |
| Bookend | Bookend | 63.6 | **84.5** | 94.0 | **52.1** | **46.5** |
| Bookend+ | None | 58.1 | 83.9 | 95.1 | 52.3 | 44.5 |
| Bookend+ | Bookend | 66.6 | 85.8 | 94.5 | 56.2 | 49.9 |
| Bookend+ | Bookend (stringent) | 77.8 | 87.8 | 94.8 | 58.3 | 54.1 |

+ Hybrid assembly with mESC CAGE and 3P-seq data

### SUPPORTING NOTES

#### The End Labeled Read file format

End Labeled Read (ELR) files are defined in two parts: a header that builds an index of reference chromosomes (#C) and read sources (#S), and a 7-column body with the following contents:

1. Chromosome index [int]
2. Alignment start position [int]
3. Alignment length (including gaps) [int]
4. Strand {+, -, .}
5. ELCIGAR string describing alignment labels and gaps [str]
6. Source of read [int]
7. Weight of all reads matching this description [float]

ELCIGAR strings describe the position and labels of all aligned segments of a read, patterned off of the BAM/SAM format CIGAR strings but with additional End Label information. They are strings of Character/Number pairs with one trailing character ($[CN]_x C$), where C is a label and N is a numeric distance on the genome. Each label annotates the end of the associated span as one of the following:

| | |
|---|---|
| S | Start Tag (RNA $5'$ end) |
| C | Capped Tag ($5'$ end with untemplated G) |
| E | End Tag (RNA $3'$ end) |
| D | Splice junction donor |
| A | Splice junction acceptor |
| . | Unspecified gap/end |

For example, the ELCIGAR for a 50bp paired-end read of a 185nt cDNA fragment with no introns or end labels would be ".50.85.50.". A full-length transcript with the ELCIGAR "$S256D800A128D800A512E$" has 3 exons of 256, 128, and 512nt, respectively, and 2 introns that are both 800nt.

Short indels and mismatches are not recorded in ELCIGAR strings, but the number of alignment errors within each exonic region (between adjacent SD, CD, AD, and AE pairs) is tallied. If this tally exceeds the user-specified error rate as a proportion of the number of aligned bases in that exon, then it is removed from the ELCIGAR and the surrounding exons are bridged with an unspecified gap (.. pair). This setting prevents the use of splice junctions from especially error-prone alignments, which can be common in some long-read sequencing protocols.

During assembly, the "weight" column will be used to determine read coverage depth for exonic frags and for end positions. However, if a Start Tag, Capped Tag, or End Tag is lowercase (s, c, e), `bookend assemble` will treat these ends as having a weight of 1 instead of the read weight. This is beneficial for partial assembly of sparsely-labeled samples.

#### Assembly Algorithms

This section lays out pseudocode for the essential elements of end-guided assembly. Pseudocode is written in a "Pythonic" way, e.g. indices and ranges are 0-indexed open.

*NOTE:* Some basic mathematical notation is used.

| | |
|---|---|
| $\sum$ | Sum |
| $\|A\|$ | Number of elements in A |
| $a : b$ | Iterate from a to b |
| $a\|b$ | a or b |
| $a \in A$ | a is in set A |
| $A \bigcup B$ | Union of sets A and B |
| $A \subset B$ | A is a subset of B |
| $A \setminus B$ | The subset of A not shared with B |

First, the input reads must be grouped into Chunks. To assemble each chunk, the branchpoints must first be generated, a list of all start clusters, end clusters, splice donors, and splice acceptors on each strand. Splice junctions are cataloged first and rare junctions are removed. Then, Tag Clustering is performed to generate two strand-specific lists of Start Clusters and Tag Clusters. Each adjacent pair of branchpoints demarcates a "frag", and the Membership of *reads* is calculated in which each read either includes (1), excludes (-1) or does not overlap with (0) each frag. Rows with identical memberships are combined to produce the Membership Matrix, and from here the Overlap Matrix can be calculated. The Overlap Matrix is further simplified by collapsing Linear Chains, after which the Overlap Graph can be defined. The weights of shorter reads fully contained within one or more longer reads are redistributed via the Resolve Containment algorithm.

Assembly is carried out on the Overlap Graph using the Greedy Paths algorithm, where a set of optimal paths through the Overlap Graph is produced. After addition of each path, the weights of all reads are assigned proportionally to each path. Finally, the optimal set of paths is filtered to remove incomplete and low-confidence models, and the remaining paths are output as assembled transcripts.

The assembly environment is built around the *RNAseqMapping* object, which acts as a general container for RNA-seq data or transcript models, and possesses a number of methods for conversion from/to various RNA-seq file formats (BED, BED12, BAM/SAM, ELR, GTF/GFF3). The data contained in each *RNAseqMapping* object includes:

*RNAseqMapping* attributes:

| name | data type | description |
|---|---|---|
| chrom | *int* | Chromosome number |
| source | *int* | Source number |
| strand | $\{1, -1, 0\}$ | Alignment strand (forward, reverse, unstranded) |
| ranges | *list* | Ordered list of (left,right) aligned segments |
| splice | *list* | List of bools $\|ranges\| - 1$; is the gap an intron? |
| weight | *float* | Abundance (counts) of the Object |
| s_tag | *bool* | Object contains a Start Tag? |
| capped | *bool* | Object contains a Capped Tag? |
| e_tag | *bool* | Object contains an End Tag? |
| complete | *bool* | Object is a full-length gapless transcript |
| is_reference | *bool* | Object came from a reference annotation file |
| condensed | *bool* | Object is a product of `bookend condense` |
| s_len | *int* | Length of trimmed Start Tag |
| e_len | *int* | Length of trimmed End Tag |
| attributes | *dict* | Container for additional $key : value$ information |

##### Generate Chunks

Assembly begins by building coherent chunks of overlapping reads that could putatively be considered "genes" or "loci". Each time there is a gap greater than the specified `--maxgap` between the rightmost edge of the reads in the chunk and the left edge of the next read, a chunk is completed. Within this chunk, false positive splice junctions may skip over gaps. To filter spurious spliced reads, read coverage (sum of read.weight) is calculated for each base in the chunk, and the weight of each splice junction is compared to the coverage of its flanking positions to discard all reads with sufficiently rare junctions.

**Algorithm 1:** GenerateChunks

**Data:** *Reads* = a sorted list of *RNAseqMapping* objects
$g$ = the maximum tolerable gap length
$p$ = the minimum proportion of coverage needed to treat a splice junction as valid
**Result:** An ordered set of *Chunks*, each a subset of *Reads* separated by a gap $> g$

```
begin
    chunk ← An empty list
    chrom ← −1
    for read ∈ Reads do
        if chunk is empty then
            spanRight ← read.right
        else if read.chrom ≠ chrom or read.left > spanRight + g then
            junctions ← Unique set of spliced positions in chunk
            junctionWeights ← Dict of ∑ r.weight for r ∈ chunk which contain each j ∈ junctions
            cov ← ∑ r.weight for all r ∈ chunk overlapping each genomic position in chunk
            badJunctions ← All j ∈ junctions where
              junctionWeights[j] < p ∗ max(cov[j.left], cov[j.right])
            chunk ← Subset of r ∈ chunk that do not contain badJunctions
            breaks ← Runs > g where no reads in chunk overlap
            for b ∈ breaks do
                subchunk ← Subset of r ∈ chunk where r.right < b
                yield subchunk
            chunk ← [read] /* Start a new chunk containing only the current read */
            spanRight ← read.right
        else
            spanRight ← max(spanRight, read.right)
        chunk.append(read)
    yield chunk
```

##### Tag Clustering

From an array of Start Tag or End Tag positions (the $5'$ or $3'$ terminus of all tagged reads, respectively), Tag Clusters are determined independently on each strand. Clusters are not allowed to span same-stranded splice junctions, which could yield malformed transcript models with termini (defined as the peak signal position within a Tag Cluster) internal to a splice site. For the same reason, same-stranded Start Clusters and End Clusters that overlap are further subdivided in a way that prevents malformed transcripts with a $3'$ end upstream of the $5'$ end (`_assembly_utils.Locus.resolve_overlapping_ends`, not shown).

**Algorithm 2:** GenerateTagClusters

```
Data: counts = Array of tag weights of one type (Start, End) and one strand (+,-) by genomic position
coverage = Array of weights of all overlapping reads by genomic position
junctions = Array of splice donor and acceptor site positions on the same strand as counts
strandRatio = Array by genomic position of inferred proportion of overlapping reads that align to the
input strand [0-1]
overhang = Minimum distance allowed between a splice site and a Tag Cluster
g = the maximum tolerable gap between tags of the same cluster
p = minimum proportion of tag weights to use as signal threshold
Result: List of TagCluster objects that record left, right, and most abundant positions
begin
    values ← counts^2 * strandRatio/coverage where counts ≠ 0
    prohibited ← iterator({junctions ± overhang, −1})
    threshold ← p * ∑ values
    passPositions ← positions where values > threshold
    TagClusters ← An empty list
    boundary ← −1
    cluster ← TagCluster(passPositions[0])
    while prohibited exists and boundary < cluster.left do
        boundary ← next(prohibited)
    for p ∈ order(passPositions[1 :], key = values[passPositions[1 :]]) do
        crossedBoundary ← boundary > −1 and p > boundary
        if crossedBoundary or p − g > cluster.right then
            TagClusters.append(cluster)
            cluster ← TagCluster(p)
            if crossedBoundary then
                while prohibited exists and boundary < p do
                    boundary ← next(prohibited)
        else
            cluster.add(p)
    TagClusters.append(cluster)
    return TagClusters
```

##### Calculate Membership Matrix

When the full set of splice junctions and clusters has been determined, the locus can be divided into $frags$, defined as a $(left, right)$ range of positions between two adjacent "branchpoints": a splice donor or acceptor site, the upstream-most position of a Start Tag Cluster, or the downstream-most position of an End Tag Cluster. The "membership" of a $read$ is an integer array of length $|frags| + 4$ that records whether it overlaps (1), does not overlap (0), or excludes (-1) the $frag$. The 4 extra integers record presence/absence/exclusion of the 4 possible Tag types: Start+, End+, Start−, End−. An exclusion means that this $read$ could not be part of any transcript that includes that $frag$; either the $read$ passes over $frag$ via an intron, or the $read$ has a Start Tag or End Tag that would terminate outside the $frag$ range. Reads with the same membership arrays are identical for the purposes of assembly, so they are collapsed into a single $element$. The Membership Matrix records the membership arrays of all $elements$ in a matrix of shape $|elements| \times |frags|$. The summed weights ($\sum readBases / \sum memberBases$) of each $element$ are recorded separately in a Weight Matrix of shape $|elements| \times |sources|$.

**Algorithm 3:** CalculateMembershipMatrix

**Data:** $Reads$ = a sorted list of $RNAseqMapping$ objects
$frags$ = Ordered list of $(left, right)$ positions of all adjacent non-overlapping branchpoints
$TagClusters$ = Collection of clusters defined by **GenerateTagClusters**
**Result:** $Membership$ : A $|elements| \times |frags| + 4$ matrix associating each $frag$ and Tag type to each unique $element$
$Weights$ : A $|elements| \times |sources|$ float matrix recording summed weights by source of each $element$
**begin**
  $Sp, Ep, Sm, Em \leftarrow |frags|, |frags|+1, |frags|+2, |frags|+3$
  $elements \leftarrow$ An empty set
  $elementWeights \leftarrow$ An empty dict
  **for** $i \in |Reads|$ **do**
    $read \leftarrow Reads[i]$
    $membership \leftarrow [0] \times (|frags| + 4)$
    **if** $read.strand = 1$ **then**
      $membership[Sp] \leftarrow int(read.s\_tag$ and $read$ starts in a Start+ Cluster)
      $membership[Ep] \leftarrow int(read.e\_tag$ and $read$ ends in an End+ Cluster)
      $membership[Sm] \leftarrow -1$
      $membership[Em] \leftarrow -1$
    **else if** $read.strand = -1$ **then**
      $membership[Sm] \leftarrow int(read.s\_tag$ and $read$ starts in a Start− Cluster)
      $membership[Em] \leftarrow int(read.e\_tag$ and $read$ ends in an End− Cluster)
      $membership[Sp] \leftarrow -1$
      $membership[Ep] \leftarrow -1$
    $previousFrag \leftarrow -1$
    **for** $j \in |read.ranges|$ **do**
      $left, right \leftarrow read.ranges[j]$
      $leftFrag \leftarrow$ which $frag$ contains $left$
      $rightFrag \leftarrow$ which $frag$ contains $right$
      $membership[leftFrag : (rightFrag + 1)] \leftarrow 1$
      **if** $j > 0$ **then**
        $gapRange \leftarrow (prevFrag + 1) : leftFrag$
        $membership[gapRange] \leftarrow$
        $$\begin{cases} -1, & \text{if } read.splice[j-1] \\ 1, & \text{if } !read.splice[j-1] \text{ and } gapRange \text{ overlaps no junctions} \\ 0, & \text{otherwise} \end{cases}$$
    $hash \leftarrow asString(membership)$
    $elements.add(hash)$
    $elementWeights[hash, read.source] \leftarrow$
     $read.weight * \sum_{r \in read.ranges}(r[1] - r[0]) / \sum_{m \in members}(frags[m][1] - frags[m][0])$
  $orderedElements \leftarrow sort(asArray(h)$ for $h \in elements)$
  $Membership \leftarrow Matrix(orderedElements)$
  $Weights \leftarrow Matrix(elementWeights[e]$ for $e \in orderedElements)$
  **return** $Membership, Weights$

##### Calculate Overlap Matrix

The Overlap Matrix ($O$) is a $|elements| \times |elements|$ square matrix that describes the relationship between each *element* pair. It is asymmetric; each pair of *elements* {a,b} has two coordinates on the matrix, $O_{ab}$ and $O_{ba}$. There are four possible overlap relationships: "does not overlap" (0), "extends" (1), "cannot extend" (-1), and "is subset of" (2). For example, if *element* b contains all members of a, then "a is subset of b" and $O_{ab} = 2$. All *elements* necessarily are subsets of themselves, so the main diagonal of $O$ is 2. "Extension" can be understood as the ability to traverse (from left to right) the *frags* of one *element* onto the *frags* of another. Examples of *element* pairs are given in the table below. The symbol set $\{-, \ , +\}$ is used for *membership* values of $\{-1, 0, 1\}$, respectively. Because alignments can contain gaps, it is possible that "a extends b" and "b extends a". Positive values in $O$ will form edges in a directed graph (the **Overlap Graph**). Negative values in $O$ can form an undirected graph (incompatibility is symmetric) that imposes limits on traversal of the Overlap Graph.

Examples of Overlap between *element* pairs

| | Membership | $O_{ab}$ | $O_{ba}$ |
|---|---|---|---|
| a<br>b | ++<br> ++ | 0 | 0 |
| a<br>b | ++<br> ++ | 1 | 0 |
| a<br>b | +++<br> ++ | 0 | 2 |
| a<br>b | ++-++<br> +++ | -1 | -1 |
| a<br>b | ++ ++<br> +++ | 1 | 1 |

**Algorithm 4:** GetOverlap

**Data:** $a$ = A single row of the Membership Matrix (integer array length $|frags| + 4$)
$b$ = A second row of the Membership Matrix
**Result:** $(O_{ab}, O_{ba})$ : Overlap relationship of $a \to b$ and $b \to a$, respectively
**begin**
  $info_a \leftarrow \sum a \neq 0$
  $info_b \leftarrow \sum b \neq 0$
  $buffer \leftarrow (False, False, False, False)$ `/* Stores` *membership* `transition states for` $(a_i, a_{i-1}, b_i, b_{i-1})$ `*/`
  $shared, ext_{ab}, ext_{ba} \leftarrow 0$
  $overlapping, in_a, in_b \leftarrow False$
  **for** $i \in 0 : |a|$ **do**
    **if** $(a_i = 1 \text{ and } b_i = -1)$ *or* $(a_i = -1 \text{ and } b_i = 1)$ **then**
      **return** $(-1, -1)$
    $shared.add(int(a_i = b_i \text{ and } a_i \neq 0))$
    $overlapping \leftarrow overlapping \text{ or } (a_i + b_i = 2)$
    $buffer \leftarrow (a_i \neq 0, buffer[0], b_i \neq 0, buffer[2])$
    $ext_{ab}.add(int(in_a \text{ and } buffer = (False, True, True, True)))$
    $ext_{ba}.add(int(in_b \text{ and } buffer = (True, True, False, True)))$
    $in_a \leftarrow ia \neq 0 \text{ and } (in_a \text{ or } a_i = 1)$
    $in_b \leftarrow ib \neq 0 \text{ and } (in_b \text{ or } b_i = 1)$
  **if** $shared \leq 0$ **then**
    **return** $(0, 0)$
  **if** $shared = info_a$ **then**
    $O_{ab} \leftarrow 2$
  **else if** $shared = info_b$ **then**
    $O_{ba} \leftarrow 2$
  **else**
    $O_{ab} \leftarrow int(overlapping \text{ and } ext_{ab} > 0)$
    $O_{ba} \leftarrow int(overlapping \text{ and } ext_{ba} > 0)$
  **return** $(O_{ab}, O_{ba})$

**Algorithm 5:** CalculateOverlapMatrix

```
Data: Membership = Output from CalculateMembershipMatrix
Strand = Array of length |elements| of alignment strands {−1, 0, 1}
Result: Overlap : Overlap Matrix, integer array of shape |elements| × |elements|
begin
    Overlap ← 0-matrix of shape |elements| × |elements|
    for a ∈ 0 : |elements| do
        for b ∈ a : |elements| do
            if b = a then
                Overlap_ab, Overlap_ba ← 2
            else
                if (Strand_a = 1 and Strand_b = −1) or (Strand_a = −1 and Strand_b = 1) then
                    Overlap_ab, Overlap_ba ← −1
                    next
            Overlap_ab, Overlap_ba ← GetOverlap(a,b)
    return Overlap
```

##### Collapse Linear Chains

When considering paths that join sets of *elements*, some relationships between *elements* are trivial, meaning that there are no branches in the path. For assembly, a non-branching set of *elements* can be treated as a single *element* that contains the sum of *membership* information in the set. To identify all non-branching *element* sets (Linear Chains), a strategy was developed to traverse the positive edges of the Overlap Matrix via Depth First Search (DFS). The *elements* are ordered by increasing information content, and all *elements* are visited by DFS. Upon postvisit, a "chain number" is assigned to the *element* based on the set of chains seen during the visit. The strategy defined below is sufficient to resolve both trivial cases and nested cases, where gapped reads create cycles in the Overlap Matrix. After labeling each *element* according to its *chain*, a new Reduced Membership Matrix ($RMM$) and Reduced Overlap Matrix ($ROM$) can be created from the set of unique *chains*, rather than all unique *elements*. The example below shows the $Membership$ and *Overlap* of a Locus before (left) and after (right) reduction. This Locus contains (1) a long linear chain, (2) a nested loop, and (3) a "long read" *element* that has complete information. The *chain* of each *element* defined by **IdentifyLinearChains** is shown in the rightmost column. The $RMM$ and $ROM$ produced by **CollapseLinearChains** are shown to the right, where *elements* that gained information from their *chain* are capitalized. It is important to note here that even though every *element* except $h$ are contained in $a$, they are assigned to a different *chain* than $a$ because, for example, *elements* $b$ through $g$ could belong to a valid path containing either $h$ or $i$, but $a$ is only compatible with $i$. If the contained *elements* were merged into a chain with $a$, then no complete path containing $h$ could exist. For *Overlap* and $ROM$, the symbol set $\{-,\ ,\circ,\bullet\}$ is used to replace $\{-1,0,1,2\}$, respectively.

| | Membership | Overlap | chain |
|---|---|---|---|
| | | abcdefghijk | |
| a | ++-+-+-++++++ | ● - | 1 |
| b | ++ | ●●○ | 5 |
| c | +-+ | ● ●○ | 5 |
| d | +-+ | ● ●○ | 5 |
| e | +-+ | ● ● ○ | 5 |
| f | + + | ● ●○ | 5 |
| g | ++ + | ● ●○○ ○ | 5 |
| h | +-+ | - ●-○ | 3 |
| i | +++ | ● -●○ | 4 |
| j | ++ | ● ●○ | 2 |
| k | ++ | ● ● | 2 |

| | RMM | ROM |
|---|---|---|
| | | aBhiJ |
| a | ++-+-+-++++++ | ● - |
| B | ++-+-+-++ ++ | ●●○○ |
| h | +-+ | - ●-○ |
| i | +++ | ● -●○ |
| J | +++ | ● ● |

**Algorithm 6:** IdentifyLinearChains

**Data:** $O$ = An adjacency list $\{a \rightarrow [b, ...]\}$ for all $a \in |elements|$ where $Overlap_{ab} > 0$
$X$ = An adjacency list $\{a \rightarrow [b, ...]\}$ for all $a \in |elements|$ where $Overlap_{ab} = -1$
$searchOrder = order(\sum e \neq 0 \text{ for } e \in elements)$
**Result:** $chains$ : List of length $|elements|$ of chain ID numbers
**begin**
- $CO \leftarrow$ An empty adjacency list
- $CX \leftarrow$ An empty adjacency list
- $vertices \leftarrow |O|$
- $visited \leftarrow$ Empty boolean array of length $vertices$
- $component \leftarrow -1$ integer array of length $vertices$
- $chainCount \leftarrow 0$
- **for** $v \in searchOrder$ **do**
  - **Visit(v)**

**method** Visit(v):
- $visited_v \leftarrow True$
- **for** $w \in O[v]$ **do**
  - **if** $not\ visited_w$ **then**
    - **Visit(w)**
- $outgroups \leftarrow unique(chain_{O[v]})$
- $outgroups.remove(-1)$
- **for** $out \in outgroups$ **do**
  - **if** $O[v] \subset CO[out]$ $and$ $X[v] = CX[out]$ **then**
    - $chain_v \leftarrow out$
    - $CO[out].add(v)$
    - **return**
- $chainCount += 1$
- $chain_v \leftarrow chainCount$
- $CO[chainCount] \leftarrow \{O[v], v]\}$
- $CX[chainCount] \leftarrow X[v]$

**Algorithm 7:** CollapseLinearChains

**Data:** $Membership$ = Output from **CalculateMembershipMatrix**
$chains$ = Output from **IdentifyLinearChains**
**Result:** $RMM$ : Reduced Membership Matrix of shape $|chains| \times |frags| + 4$
**begin**
- $RMM \leftarrow copy(Membership)$
- $chainIndices \leftarrow$ An empty list
- $chainDict \leftarrow$ An empty dict
- **for** $i \in |vertices|$ **do**
  - **if** $chain_i \in keys(chainDict)$ **then**
    - $parent \leftarrow chainDict[chain_i]$
    - $newInfo \leftarrow where(Membership[i, :] \neq 0$
    - $RMM[parent, newInfo] \leftarrow Membership[i, newInfo]$
  - **else**
    - $chainIndices.append(i)$ $chainDict[chain_i] \leftarrow i$
- **return** $RMM[chainIndices, :]$

##### Generate Overlap Graph

With matrices of $Membership$, $Overlap$, $Weights$ calculated for all *chains*, it is now possible to construct the Overlap Graph where $Node$ objects are assembled into *Paths*. A $Node$ object is constructed for each *chain*, and *Paths* across multiple *chains* can also be expressed as a $Node$:

$Node$ attributes:

| name | data type | description |
|---|---|---|
| index | *int* | Ordered ID number |
| length | *int* | Total exonic length (nucleotides) |
| IC | *int* | Information Content, $\sum info \neq 0$ |
| maxIC | *int* | $\lvert frags \rvert + 4$ |
| left | *int* | Lowest *index* of contained $Nodes$ |
| right | *int* | Highest *index* of contained $Nodes$ |
| members | *set* | $f \in frag$ where $membership_f = 1$ |
| nonmembers | *set* | $f \in frag$ where $membership_f = -1$ |
| LM | *int* | $min(members)$ |
| RM | *int* | $max(members)$ |
| strand | $\{-1, 0, 1\}$ | Alignment strand |
| bases | *float* | Total sequenced bases contained in $Node$ |
| cov | *float* | $bases/length$ |
| weights | *array* | $\lvert sources \rvert$, cov contributed by each source |
| member_weights | *array* | $\lvert frags \rvert$, estimated cov of each $frag$ |
| ingroup | *set* | $\{i \in \lvert Nodes \rvert$ where $Overlap_{i,Node.index} = 1\}$ |
| outgroup | *set* | $\{i \in \lvert Nodes \rvert$ where $Overlap_{Node.index,i} = 1\}$ |
| contains | *set* | $\{i \in \lvert Nodes \rvert$ where $Overlap_{i,Node.index} = 2\}$ |
| contained | *set* | $\{i \in \lvert Nodes \rvert$ where $Overlap_{Node.index,i} = 2\}$ |
| excludes | *set* | $\{i \in \lvert Nodes \rvert$ where $Overlap_{Node.index,i} = -1\}$ |
| includes | *set* | Set of indices of $Nodes$ included in $Path$ |
| assigned_to | *set* | Set of $Paths$ this $Node$ is part of |
| complete | *bool* | This $Node$ is a full-length, gapless transcript |
| s_tag | *bool* | This $Node$ has a Start Tag |
| e_tag | *bool* | This $Node$ has an End Tag |
| is_spliced | *bool* | This $Node$ contains at least one intron |
| has_gaps | *bool* | This $Node$ contains at least one internal gap |

The set of $Nodes$ generated for the Overlap Graph are $elementNodes$; they map 1-to-1 to the rows of the Reduced Membership Matrix. New $Node$ objects can be constructed by combining multiple $elementNodes$, and any valid set of $elementNodes$ connected by edges in the Overlap Matrix with no incompatibilities is a $Path$.

**Algorithm 8:** OverlapGraph Constructor

**Data:** $M$ = Reduced Membership Matrix, shape $\lvert chains \rvert \times \lvert frags \rvert + 4$
$O$ = Reduced Overlap Matrix, shape $\lvert chains \rvert \times \lvert chains \rvert$
$sourceWeights$ = Matrix recording sequenced $bases/length$, shape $\lvert chains \rvert \times \lvert sources \rvert$
$fragWeights$ = Matrix estimating coverage by $frag$ for each *chain*, shape $\lvert chains \rvert \times \lvert frags \rvert$
**Result:** *OverlapGraph* object: Container for the lists of $elementNodes$ and $Paths$
**begin**

```
    OG ← An empty OverlapGraph object
    OG.elementNodes ← An empty list
    OG.Paths ← An empty list
    for i ∈ |chains| do
        OG.elementNodes.append(Node(M[i, :], O[i, ], sourceWeights[i, :], fragWeights[i, :])
        if OG.elementNodes_i.complete then
            OG.Paths.append(copy(OG.elementNodes_i))
    OG.bases ← ∑(e.bases for e ∈ OG.elementNodes)
    return OG
```

**Resolve Containment**

When the *OverlapGraph* object is constructed, there will usually be a number of *Nodes* that are contained by other *Nodes*. A contained *Node* is a subset of one or more containers, and assembly performance is improved the weight of contained *Nodes* is passed proportionally to its containers prior to calculating *Paths*. The Resolve Containment algorithm "bubbles up" weight from contained *Nodes* in order of decreasing information content (longest first). The total weight of the Locus is preserved, but the weights of all contained elements that cannot form a *Path* incompatible with their containers are set to zero. This allows the shorter *elementNodes* to be ignored during assembly without losing their coverage information.

---

**Algorithm 9:** ResolveContainment

---

**Data:** $OG$ = Overlap Graph Object
**Result:** In-place update of $weights$ in $OG.elementNodes$
**begin**
  $zeros \leftarrow$ Set of all $OG.elementNodes$ with $cov = 0$
  $containedNodes \leftarrow$ which $|e.contained| > 0$ for $e \in OG.elementNodes$
  $resolveOrder \leftarrow order(containedNodes, key1 = IC, key2 = |contained|)$
  **for** $i \in resolveOrder$ **do**
    $element = OG.elementNodes_i$
    **if** $element.cov > 0$ **then**
      $containers \leftarrow element.contained \setminus zeros$
      $incompatible \leftarrow set(0 : |OG.elementNodes|) \setminus zeros$
      **for** $c \in containers$ **do**
        $incompatible \leftarrow incompatible \bigcap OG.elementNodes_c.excludes$
      $incompatible \leftarrow incompatible \setminus element.excludes$
      **if** $|incompatible| = 0$ **then**
        $containerCov \leftarrow$ Array of $OG.elementNodes_c.cov$ for $c \in containers$
        $totalCov \leftarrow \sum containerCov$
        $defaultProportions \leftarrow$ Array length $|containers|$ of $containerCov/totalCov$
        $proportions \leftarrow$ 0-array shape $|containers| \times |sources|$
        **for** $i \in 0 : |containers|$ **do**
          $c \leftarrow containers_i$
          $weights \leftarrow OG.elementNodes_c.weights$
          $proportions[i, :] \leftarrow defaultProportions \times weights / \sum weights$
        **for** $i \in 0 : |sources|$ **do**
          **if** $\sum proportions[:, i] = 0$ **then**
            $proportions[:, i] \leftarrow defaultProportions$
          **else**
            $proportions[:, i] \leftarrow proportions[:, i] / \sum proportions[:, i]$
        **for** $i \in 0 : |containers|$ **do**
          $c \leftarrow containers_i$
          $cNode \leftarrow OG.elementNodes_c$
          $cNode.weights.add(element.weights \times proportions[i, :])$
        $element.weights \leftarrow$ 0-array of length $|sources|$
        $zeros.add(i)$

---

##### Greedy Paths

The Overlap Graph can be understood to have a $source(s)$ and $sink(t)$, and regardless of alignment strand, all edges between $Nodes$ describe flow from left to right in genomic positions. The four Tag types connect to the $source(Start+, End-)$ and $sink(Start-, End+)$, so any complete $s \rightarrow t$ Path must contain a same-stranded pair of Start/End Tags. To find an optimal path through the OverlapGraph, search begins at the $Node$ with the greatest weight, and examines each edge to continue outward in a Breadth-First Search that always traverses the edge with the highest "Extension Score". This score is an estimate of the total available weight along the extending edge, counterbalanced by 3 separate penalties:

$sourceSimilarity$: [0-1] Distance between the relative source contributions of the Path and the Extension.
$variancePenalty$: [0-1] Ratio between the mean sectional coverage and max sectional coverage of the Path.
$deadEndPenalty$: [0-1] Multiplier imposed if no complete $Paths$ can extend through this edge.

When the search terminates, the resulting set of $Nodes$ is combined to yield an "Optimal Path", and is stored in a list of $Paths$. Assembly of the locus is complete when less than `--min_proportion` of the reads are unassigned or the same path is generated twice.

**Algorithm 10:** ExtensionScore

```
Data: Path = A Node object representing an incomplete set of elementNodes
extension = A candidate list of elementNodes to add to Path
p = minimum proportion
Result: score : Float that evaluates the extension to Path (higher is score better)
begin
    extensionWeights ← 0-array of length |frags|
    sourceProportions ← 0-array of length |sources|
    newFrags ← An empty set
    for node ∈ extension do
        newFrags.add(frags in node and not in Path)
        newProportions ← Array of length |sources|,
            { 1-array if node is unassigned
            { otherwise Path.weights/(Path.weights + ∑(p.weights for p ∈ node.assigned_to))
        available ← ∑ newProportions/|newProportions|
        if available < p then
            available ← 0
        extensionWeights.add(available ∗ node.member_weights)
        sourceProportions.add(newProportions/|extension|)
    extensionCov ← max(extensionWeights[newFrags])
    combinedWeights ← Path.member_weights + extensionWeights
    pathProportions ← Path.weights/|Path.weights|
    sourceSimilarity ← .5 ∗ (2 − ∑(abs(pathProportions − sourceProportions)))
    variancePenalty ← mean(combinedWeights)/max(combinedWeights)
    deadEndPenalty ← { 0 if s or t are unreachable from Path+extension
                     { 1 otherwise
    score ← extensionCov ∗ sourceSimilarity ∗ variancePenalty ∗ deadEndPenalty
    return score
```

Using default arguments for assembly, $deadEndPenalty$ is absolute and yields a $score$ of 0 for any $extension$ to a $Path$ that cannot be part of an $s \rightarrow t$ Path. If assembly is run with the argument `--allow_incomplete`, the penalty is instead a multiplier of .1 if $s$ is unreachable, and a second .1 multiplier if $t$ is unreachable. Complete $Paths$ are still heavily favored, but a $Path$ will nonetheless be produced if $s \rightarrow t$ Paths do not exist or are extremely poor. Each step of the Greedy Paths algorithm begins by generating a set of possible extensions. Each $extension$ within $extensions$ is a set of mutually compatible $elementNodes$ that can extend the given $Path$ exactly one step to the left and/or right. Each $extension$ is composed of an ingroup (a node with an edge to $Path$) and an outgroup (a node that $Path$ has an edge to), and any nodes that may be contained by the union of the ingroup, outgroup, and $Path$. The Greedy Path algorithm begins at the heaviest unassigned $elementNode$, and each extension step lengthens the $Path$ toward both the source and sink.

**Algorithm 11:** GenerateExtensions

```
Data: OG = Overlap Graph object
Path = An incomplete merged set of elementNodes
Result: extensions : A list of candidate sets of elementNodes to extend Path
begin
    extensions ← An empty list of sets
    ingroup ← Path.ingroup ⋃ Path.contained
    outgroup ← Path.outgroup ⋃ Path.contained
    freeNodes ← Path.contains \ Path.includes
    if |freeNodes| > 0 then
        Path.extend(freeNodes)
        ingroup ← Path.ingroup
        outgroup ← Path.outgroup
    if |ingroup \ Path.contained| > 0 then
        if |outgroup \ Path.contained| > 0 then
            pairs ← List of all (i, o) pairs for i ∈ ingroup, o ∈ outgroup if Overlap[i, o] > −1
        else
            pairs ← List (i, Path.index) for i ∈ ingroup
    else
        pairs ← List (Path.index, o) for o ∈ outgroup
    for (in, out) ∈ pairs do
        e_in ← OG.elementNodes_in
        e_out ← OG.elementNodes_out
        contained ← e_in.outgroup ⋃ e_out.ingroup ⋃ e_in.contains ⋃ e_out.contains
        /* Filter 1:  All elements already included or excluded in the extension       */
        exclude ← e_in.includes|Path.includes|e_out.includes|e_in.excludes|Path.excludes|e_out.excludes
        contained ← contained \ exclude
        /* Filter 2:  All elements that add information not contained in the extension */
        stranded ← e_in.strand ≠ 0 or e_out.strand ≠ 0 or Path.strand ≠ 0
        extMembers ← e_in.members ⋃ Path.members ⋃ e_out.members
        extNonmembers ← e_in.nonmembers ⋃ Path.nonmembers ⋃ e_out.nonmembers
        exclude ← Set of contained elements if not contained.members ⊂ extMembers or not
          contained.nonmembers ⊂ extNonmembers
        contained.add(in, out)
        contained ← contained \ exclude
        extensions.add(contained)
    return extensions
```

**Algorithm 12:** GreedyPaths

```
Data: OG = Overlap Graph object
p = minimum proportion
Result: Path : The highest-scoring complete s → t path through the heaviest unassigned elementNode
begin
    Path ← node with max(node.cov for node ∈ OG.elementNodes) where |node.assignments| = 0
    Path.extend(Path.contains)
    extensions ← GenerateExtensions(Path)
    while |extensions| > 0 do
        if |extensions| = 1 then
            Path.extend(extensions_0)
        else
            bestExtension ← ext ∈ extensions with max(ExtensionScore(Path, ext))
            Path.extend(bestExtension)
        extensions ← GenerateExtensions(Path)
    return Path
```

##### Assign Weights to Paths

If an OverlapGraph has one or more $Paths$, the weights of individual $Nodes$ must be assigned proportionally to the $Paths$ of which they are parts. A $Node$ assigned to a single $Path$ assigns all its weight to that $Path$, but if multiple overlapping $Paths$ exist, weight must be assigned proportionally to each. The estimated weight of each $Path$ is calculated by using the previous $Path$ weights as priors. $Nodes$ are resolved in increasing order of number of assignments, and their weight is added proportionally to the priors of all assigned $Paths$.

**Algorithm 13:** AssignWeightsToPaths

**Data:** $OG$ = Overlap Graph object
**Result:** In-place update of $weights$ in $OG.Paths$
**begin**
- $priors \leftarrow$ 0-array of shape $|OG.Paths| \times |sources|$
- **for** $i \in 0 : |OG.paths|$ **do**
  - $priors[i, :] \leftarrow OG.Paths_i.weights$
  - $OG.Paths_i.weights \leftarrow 0 - arrayoflength|sources|$
- $pathCovs \leftarrow \sum_{j=0}^{|sources|} priors_{ij}$
- $sampleCovs \leftarrow \sum_{i=0}^{|OG.Paths|} priors_{ij}$
- $pathProportions \leftarrow pathCovs / \sum pathCovs$
- $proportions \leftarrow$ Array shape $|OG.Paths| \times |sources|$ column-filled with $pathCovs$
- **for** $i$ $where$ $sampleCovs > 0$ **do**
  - $proportions[:, i] \leftarrow priors[:, i]/sampleCovs_i$
- **for** $i \in order(OG.assignments)$ **do**
  - $element \leftarrow OG.elementNodes_i$
  - **if** $OG.assignments_i = 1$ **then**
    - $OG.Paths_{element.assigned\_to}.weight.add(element.weights)$
  - **else if** $OG.assignments_i > 1$ **then**
    - `/* Assigned paths must compete for` $element.weight$ `*/`
    - $assigned \leftarrow element.assigned\_to$
    - $assignedProportions \leftarrow proportions[assigned, :]$
    - **for** $jin0 : |assigned|$ **do**
      - $Path \leftarrow OG.Paths_{assigned_j}$
      - $Path.weight.add(element.weight * element.length * assignedProportions[j, :])$

**Path Filtering**

For a signal threshold $p$, assembly is complete after $> (1-p)$ bases in the Overlap Graph have been assigned to $Paths$. However, due to incomplete or error-prone data, the optimal set of $Paths$ may still contain a number of errors that should be removed prior to exporting the remaining $Paths$ and transcript models. Five types of possible error are identified in the order listed below and are either removed or subject to more stringent filters.

Incomplete assemblies: Any $Path$ that has gaps or is missing a Start or End Tag.
Fused assemblies: A $Path$ that fully contains two non-overlapping $Paths$.
Truncated assemblies: A $Path$ fully contained within a longer path.
Retained introns: An overlapping $Path$ has a superset of introns.
Minimum isoform proportion: A $Path$ is assigned $< p$ of the summed bases of overlapping $Paths$

The definition of a complete $Path$ is built into the $Node$ object, but the other filters need to be calculated with reference to other $Paths$ in the Overlap Graph. Each filter defines a subset of $Paths$ to keep, and in the $FilterPaths$ method weights are reassigned after each filter because the total number of $Paths$ may have changed.

**Algorithm 14:** FilterFusions

```
Data: OG = Overlap Graph object
Result: filteredPaths : Subset of OG.Paths that passed the filter
begin
    badPaths ← Empty boolean array length |OG.Paths|
    for i ∈ |OG.Paths| do
        p1 ← OG.Paths_i
        containedRanges ← An empty list
        for j ∈ |OG.Paths| ≠ i do
            p2 ← OG.Paths_j
            if p2.members ⊂ p1.members and p2.weight ≥ p1.weight then
                containedRanges.append((p2.LM, p2.RM))
        for c1 ∈ containedRanges do
            for c2 ∈ containedRanges do
                if c1_0 > c2_1 or c2_0 > c1_1 then
                    badPaths_i ← True
    filteredPaths ← OG.Paths[!badPaths]
    return filteredPaths
```

**Algorithm 15:** FilterTruncations

```
Data: OG = Overlap Graph object
Result: filteredPaths : Subset of OG.Paths that passed the filter
begin
    badPaths ← Empty boolean array length |OG.Paths|
    for i ∈ |OG.Paths| do
        containerCov ← 0
        p1 ← OG.Paths_i
        for j ∈ |OG.Paths| do
            p2 ← OG.Paths_j
            if p1 overlaps p2 and p1.members ⊂ p2.members then
                containerCov.add(p2.cov)
        if containerCov > 0 and p1.cov < p1.cov + containerCov then
            badPaths_i ← True
    filteredPaths ← OG.Paths[!badPaths]
    return filteredPaths
```

**Algorithm 16:** FilterRetainedIntrons

```
Data: OG = Overlap Graph object
intronFilter = Minimum proportion of coverage to keep a retained intron
Result: filteredPaths : Subset of OG.Paths that passed the filter
begin
    badPaths ← Empty boolean array length |OG.Paths|
    for i ∈ |OG.Paths| do
        containerCov ← 0
        p1 ← OG.Paths_i
        for j ∈ |OG.Paths| do
            p2 ← OG.Paths_j
            if p1.LM = p2.LM and p1.RM = p2.RM then
                if introns(p1) ⊂ introns(p2) then
                    containerCov.add(p2.cov)
        if containerCov > 0 and p1.cov < intronFilter ∗ (p1.cov + containerCov) then
            badPaths_i ← True
    filteredPaths ← OG.Paths[!badPaths]
    return filteredPaths
```

**Algorithm 17:** FilterMinimumProportion

```
Data: OG = Overlap Graph object
p = Minimum proportion
Result: filteredPaths : Subset of OG.Paths that passed the filter
begin
    badPaths ← Empty boolean array length |OG.Paths|
    for i ∈ |OG.Paths| do
        overlapCov ← 0
        p1 ← OG.Paths_i
        for j ∈ |OG.Paths| do
            p2 ← OG.Paths_j
            if |p1.members ⋂ p2.members| ≥ .5 ∗ min(|p1.members|, |p2.members|) then
                overlapCov.add(p2.cov)
        if overlapCov > 0 and p1.cov < p ∗ (p1.cov + overlapCov) then
            badPaths_i ← True
    filteredPaths ← OG.Paths[!badPaths]
    return filteredPaths
```

**Algorithm 18:** FilterPaths

```
Data: OG = Overlap Graph object with a complete set of Optimal Paths
Result: In-place update of OG.Paths to retain only unfiltered Paths
begin
    OG.Paths ← p ∈ OG.Paths if p.complete
    AssignWeightsToPaths(OG)
    OG.Paths ← FilterFusions(OG)
    AssignWeightsToPaths(OG)
    OG.Paths ← FilterTruncations(OG)
    AssignWeightsToPaths(OG)
    OG.Paths ← FilterRetainedIntrons(OG)
    AssignWeightsToPaths(OG)
    OG.Paths ← FilterMinimumProportion(OG)
    AssignWeightsToPaths(OG)
```
